# Subventricular zone cytogenesis provides trophic support for neural repair

**DOI:** 10.1101/2022.06.14.496078

**Authors:** Michael R. Williamson, Stephanie P. Le, Ronald L. Franzen, Nicole A. Donlan, Jill L. Rosow, Andrew K. Dunn, Theresa A. Jones, Michael R. Drew

## Abstract

Stroke enhances proliferation of neural precursor cells within the subventricular zone (SVZ) and induces ectopic migration of newborn cells towards the site of injury. Here we characterize the identity of cells arising from the SVZ after stroke and provide insight into their function by uncovering a mechanism through which they facilitate neural repair and functional recovery. Using genetic lineage tracing, we show that SVZ-derived cells that migrate towards stroke-induced cortical lesions in mice are predominantly undifferentiated precursors, suggesting that the main function of post-injury cytogenesis is not cell replacement. We find that SVZ-derived cells are a unique cellular source of trophic factors that instruct neural repair. Chemogenetic ablation of neural precursor cells or conditional knockout of VEGF in the adult neural stem cell lineage impairs neuronal and vascular reparative responses and worsens functional recovery after stroke. In addition, normal aging markedly diminishes the cytogenic response to stroke, resulting in worse functional recovery. Therapeutic replacement of VEGF in peri-infarct cortex is sufficient to induce neural repair and functional recovery in mice with arrested cytogenesis. These findings indicate that the SVZ cytogenic response following brain injury is a source of trophic support that drives neural repair and recovery.

## Introduction

Limited recovery of function occurs after damage to the central nervous system. Consequently, stroke and other forms of brain injury often cause long-lasting disabilities. Following stroke, remodeling of residual tissue surrounding the site of injury is thought to underlie recovery. For example, extensive plasticity of neural circuits and blood vessels occurs in peri-infarct regions and these processes are associated with functional improvement (Brown et al., 2007; Clark et al., 2019; Tennant et al., 2017; Williamson et al., 2020). Repair processes are mediated by interactions across disparate cell types (Brown et al., 2007; Clark et al., 2019; Joy et al., 2019; Kim et al., 2018; Williamson et al., 2021). However, the intercellular interactions that orchestrate repair and recovery remain to be completely defined. A more complete understanding of the mechanisms that govern neural repair could inform development of new treatment strategies.

Cytogenesis, the formation of new cells, is limited in the adult mammalian brain. The subventricular zone (SVZ) is one of a small number of regions that contains multipotent neural stem and progenitor cells (collectively referred to here as precursors) that generate new neurons and glia in adulthood (Doetsch et al., 1999; Garcia et al., 2004). Normally, the predominant progeny arising from the SVZ are new neurons that migrate towards the olfactory bulb and integrate into existing olfactory circuitry. However, injuries such as stroke markedly increase SVZ precursor proliferation and induce ectopic migration of SVZ-derived cells towards the site of injury (Arvidsson et al., 2002; Lagace, 2012; Li et al., 2010; Ohab et al., 2006; Parent et al., 2002; Williamson et al., 2019). Past studies of this process after brain injury have largely focused on neurogenesis — the formation of new neurons and their localization in peri-infarct regions (Arvidsson et al., 2002; Parent et al., 2002). In general, experimental manipulations that increase post-stroke neurogenesis are associated with enhanced functional recovery (Lagace, 2012). The prevailing view has been that cell replacement, especially neuron replacement, by SVZ precursors allows for partial brain regeneration and consequently improved function. However, recent findings indicate that new neurons poorly integrate into existing circuits in peri-infarct regions and receive little synaptic input (Kannangara et al., 2018), making their functional importance unclear (cf. Liang et al., 2019). Moreover, other studies have found that glia outnumber neurons among migrating SVZ-derived cells, but the entire population of SVZ-derived cells has yet to be comprehensively characterized (Benner et al., 2013; Faiz et al., 2015; Li et al., 2010). Overall, the identity and functional importance of new cells that arise from the SVZ after injury are not well understood.

Here we investigate the SVZ cytogenic response following cortical photothrombotic strokes in mice. Our goals were to characterize the types of cells produced by the SVZ after stroke and mechanistically understand the role of SVZ cytogenesis in stroke recovery. We use indelible lineage tracing to phenotype SVZ-derived cells that migrate towards the site of injury. Unexpectedly, we find that the majority of these cells are undifferentiated precursors, suggesting that cell replacement is limited. Reducing cytogenesis impairs motor recovery after stroke, at least in part due to deficits in neuronal and vascular plasticity. With gain- and loss-of-function manipulations, we show that VEGF produced by SVZ-derived cells drives repair and functional recovery. These findings identify SVZ cytogenesis as a source of trophic support that facilitates neural repair. Thus, our study demonstrates a mechanism other than cell replacement by which endogenous neural precursor cells contribute to repair and recovery in the injured central nervous system.

## Results

### Cells arising from the subventricular zone after stroke are predominantly quiescent precursors and astrocytes

We used genetic lineage tracing to characterize the SVZ cytogenic response to stroke.

Young adult (3-6 months old) *Nestin-CreER*; Ai14 mice were injected with tamoxifen to induce indelible tdTomato expression in neural stem cells and their progeny (Figure 1A, B) (Benner et al., 2013; Li et al., 2010). Four weeks later, photothrombotic cortical infarcts were induced, and tissue was collected two weeks post-stroke, which is a time when large numbers of SVZ-derived cell are localized to peri-infarct cortex, substantial neural repair is ongoing, and functional improvement is incomplete (Kannangara et al., 2018; Kim et al., 2018; Williamson et al., 2021). While no cortical migration was seen in the absence of injury, unilateral strokes induced a profound migration of tdTomato^+^ cells from the SVZ into peri-infarct cortex (Figure 1C, D). We immunostained tissue to examine expression of an array of differentiation stage-specific and proliferation-associated proteins in lineage-traced cells (Figure 1E-Q, Supplemental Figure 1, Supplemental Figure 2).

**Figure 1.**
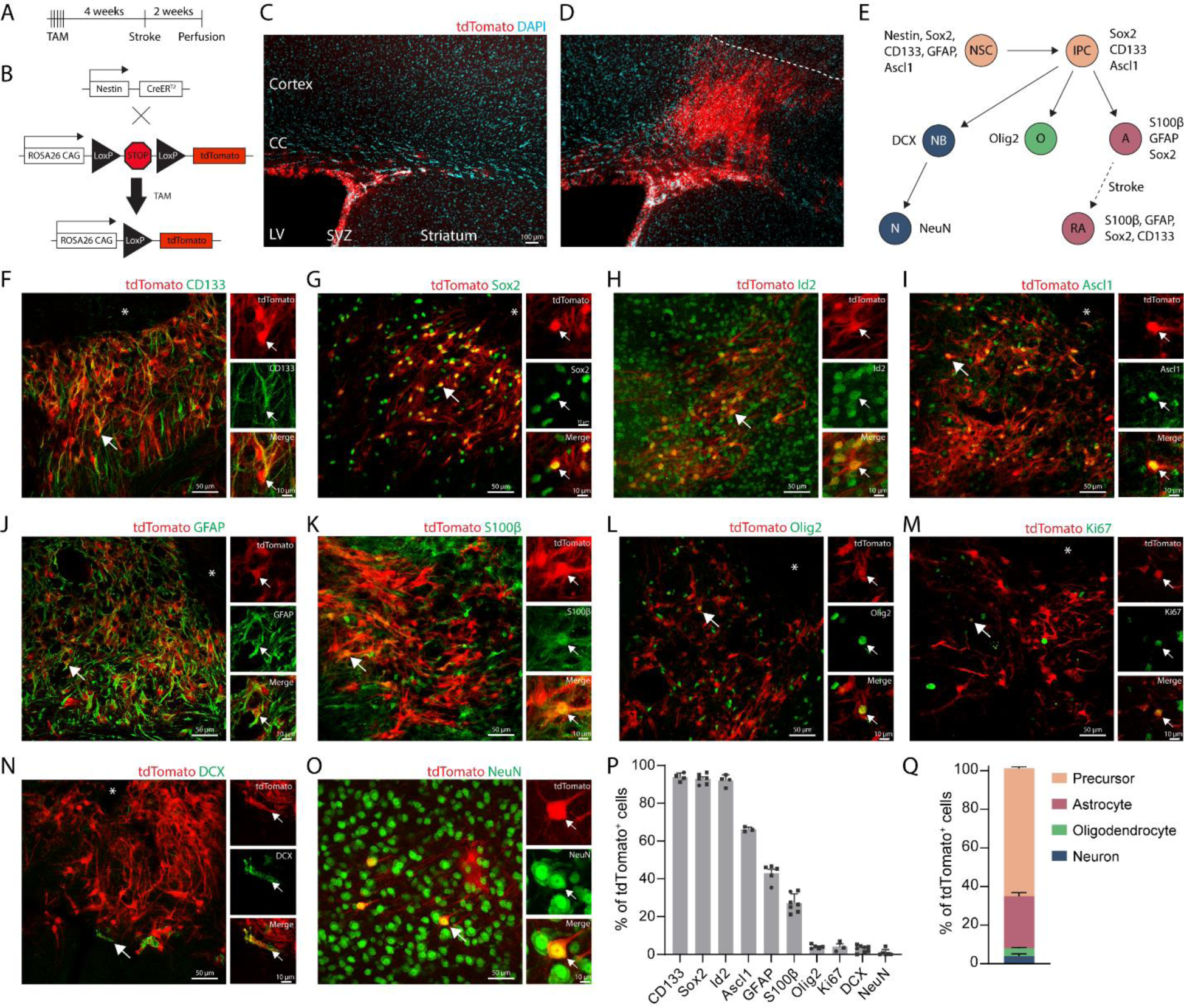
SVZ-derived cells are predominantly undifferentiated precursors and astrocytes. A) Experimental timeline for lineage tracing neural stem cell progeny after stroke. Tissue was collected two weeks post-stroke. B) Schematic of inducible, neural stem cell-specific lineage tracing system in Nestin-CreER; Ai14 mice. Tamoxifen (TAM) administration induces tdTomato expression in neural stem cells and their progeny. C) Image of tdTomato-expressing cells in the subventricular zone in the absence of injury. Note the lack of cortical migration of tdTomato^+^ cells. CC, corpus callosum; LV, lateral ventricle; SVZ, subventricular zone. D) Substantial migration of tdTomato^+^ cells towards the infarct after cortical photothrombotic stroke (dashed line indicates the approximate infarct border). E) Schematic of differentiation stages as defined by marker expression. Neural stem cells (NSC) produce intermediate progenitor cells (IPC), which give rise to cells of the three major neural lineages: neurons (NB, neuroblast; N, neuron), oligodendrocytes (O), and astrocytes (A). After stroke, reactive astrocytes (RA) re-express some neural stem cell markers. F-O) Representative confocal images of tdTomato^+^ cells and immunostaining for lineage and functional markers. Images are from peri-infarct cortex two weeks post-stroke. Asterisks indicate the lesion core. Most tdTomato^+^ cells (precursors and astrocytes) expressed the stem cell-associated markers CD133 (F) and Sox2 (G), and the quiescence marker Id2 (H). Id2^+^ tdTomato^-^ cells include mature resident cortical cells. Undifferentiated precursors were uniquely identified by expression of Ascl1 (I). J) GFAP marked astrocytes and a subset of precursors. K) Differentiated astrocytes were defined by expression of S100β. L) The oligodendrocyte lineage was defined by Olig2 expression. M) Few cells expressed the proliferation marker Ki67. Neuron lineage cells, DCX^+^ (N) and NeuN^+^ (O), were the least common. P) Quantification of marker expression by tdTomato^+^ cells. Data are from n = 3-8 mice per marker, and > 100 tdTomato^+^ cells counted per mouse per marker. Q) Estimate of phenotype distribution of lineage traced cells. Precursors were defined by Ascl1 expression. Astrocytes were defined by S100β expression. Oligodendrocyte lineage was defined by Olig2 expression. Neuron lineage was defined by DCX and NeuN expression. See also Supplemental Figure 1 and Supplemental Figure 2. Data are presented as mean ± SEM. Datapoints representing males are shown as circles; datapoints representing females are shown as squares.

Unexpectedly, the majority of tdTomato^+^ cells in peri-infarct cortex expressed precursor cell-associated markers (93.8±1.1% were CD133^+^, 92.9±1.1% were Sox2^+^, and 66.2±0.7% were Ascl1^+^). 27.1±1.9% of tdTomato^+^ cells were differentiated astrocytes based on expression of S100β, which defines astrocyte maturation and loss of multipotency (Lattke et al., 2021; Raponi et al., 2007) and is not expressed by SVZ precursors (Codega et al., 2014). Astrocyte reactivity is associated with re-expression of some precursor cell-associated proteins, including CD133 and Sox2 (Götz et al., 2015; Robel et al., 2011), but reactive astrocytes do not express Ascl1 (Magnusson et al., 2014; Zamboni et al., 2020). Thus, the Ascl1^+^ subpopulation defined undifferentiated precursors, the S100β^+^ subpopulation defined astrocytes, and CD133/Sox2 labeled both subpopulations (Figure 1E). Lineage-traced cells were largely quiescent as defined by expression of the quiescence marker Id2 in 92.1±1.5% of cells (Llorens-Bobadilla et al., 2015), and rare expression of the proliferation marker Ki67 (4.1±1.5%). Oligodendrocyte-lineage (Olig2^+^, 4.1±0.5%) and neuron-lineage cells (DCX^+^, 3.0±0.6%, and NeuN^+^, 0.9±0.7%; (also see Supplemental Figure 2)) made up the remainder of lineage-traced cells. We corroborated these phenotyping results with parallel experiments in *Ascl1-CreER*; Ai14 mice, in which a subset of neural stem and progenitor cells are lineage-traced (Liang et al., 2019) (Supplemental Figure 1). We also confirmed that few new neurons are formed up to 6 weeks post-stroke using *Nestin- CreER*; *Sun1-sfGFP^fl^* mice in which neural stem cells and their progeny were labeled with a nuclear membrane-targeted fluorophore (Supplemental Figure 2). These experiments identify undifferentiated precursors as the predominant cell type produced by the SVZ in response to stroke. Since the majority of new cells that localize to peri-infarct regions remain in an undifferentiated state, cell replacement may not be the primary function of the cytogenic response.

### Chemogenetic ablation of neural stem cells impairs recovery after stroke

Having characterized the phenotype of SVZ-derived cells, we next investigated whether SVZ cytogenesis provides functional benefits for recovery after stroke. We used GFAP-TK mice to selectively ablate neural stem cells prior to stroke following an established ganciclovir (GCV) administration paradigm (Garcia et al., 2004; Swan et al., 2014) (Figure 2A-J). Delivery of GCV via subcutaneous osmotic pumps for two weeks ablates mitotic TK-expressing cells (i.e. neural stem cells), but spares non-mitotic GFAP-expressing cells, including cortical astrocytes (Garcia et al., 2004; Swan et al., 2014) (Figure 2D-F; Supplemental Figure 3A-C). SVZ cytogenesis, as measured by the density of SVZ DCX^+^ and Ki67^+^ cells, was substantially reduced in GFAP- TK+GCV mice relative to littermate controls, which included wildtype mice given saline or GCV and GFAP-TK mice given saline (Figure 2G-J).

**Figure 2.**
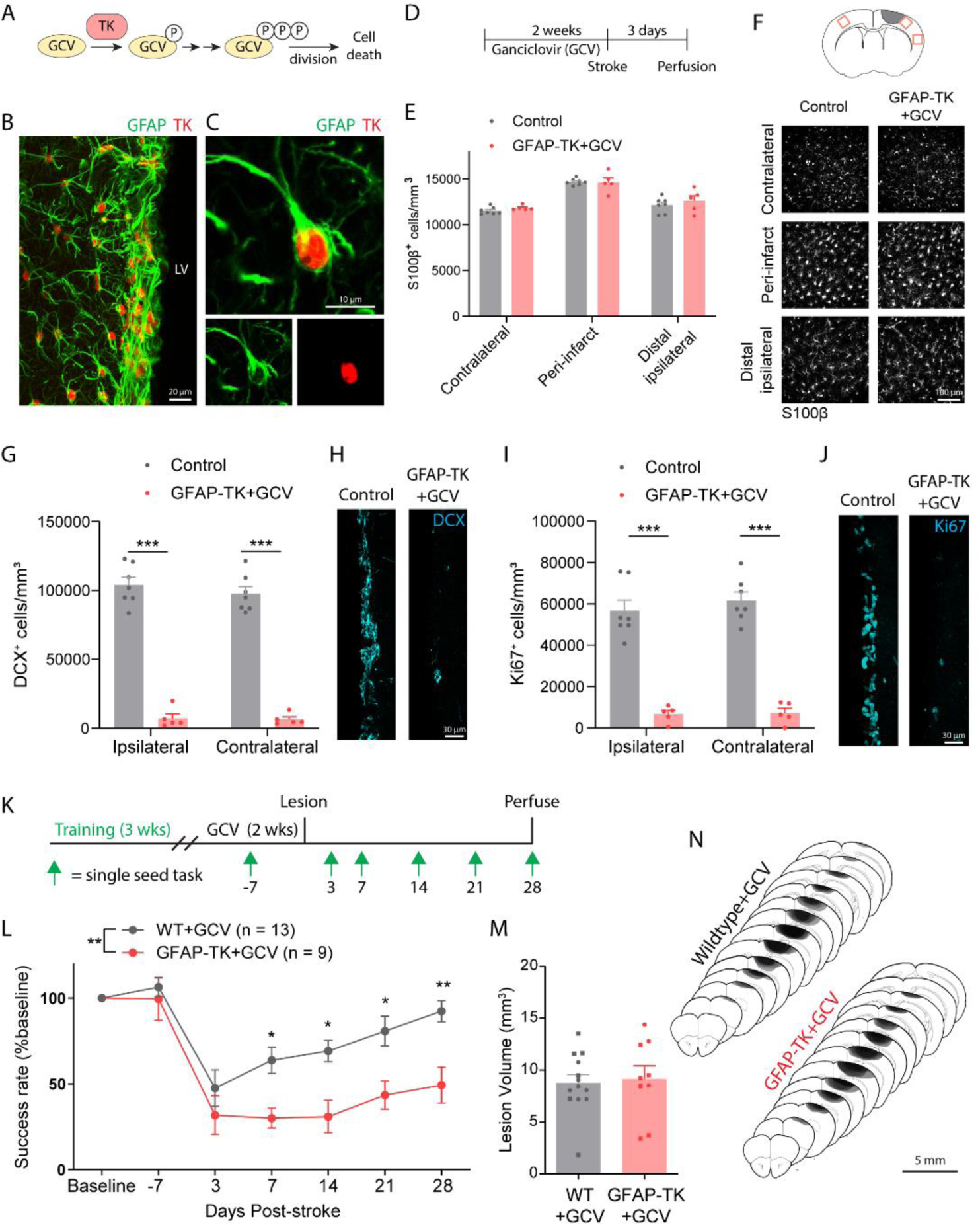
Ablation of neural stem cells worsens recovery after stroke. A) Schematic illustrating the thymidine kinase (TK)/ganciclovir (GCV) system for conditional ablation of proliferating cells. B) Confocal image showing TK expression in GFAP^+^ cells of the subventricular zone in a GFAP-TK mouse. LV, lateral ventricle. C) High-resolution image of a TK-expressing SVZ stem cell. D) Experimental timeline for assessing the specificity and effectiveness of arresting SVZ cytogenesis with GFAP-TK mice (E-J). GCV or saline was delivered for 14 days via osmotic pump. Tissue was collected 3 days after stroke. n = 7 control (wildtype mice given GCV or GFAP-TK mice given saline), n = 5 GFAP-TK+GCV. E) Parenchymal astrocytes were not depleted in this paradigm. t(10) < 1.53, p ≥ 0.157, t tests comparing groups for each region. F) Representative images of S100β^+^ cells from three cortical regions (see diagram at top) show lack of astrocyte ablation. G-J) GFAP-TK mice permit conditional arrest of cytogenesis. The number of SVZ DCX^+^ (G, H) and Ki67^+^ cells (I, J) was significantly reduced in GFAP-TK mice given GCV relative to controls. ***t(10) ≥ 7.99, p < 0.001, t tests. K) Experimental design for assessing motor recovery after photothrombotic stroke with the single seed reaching task (n = 13 wildtype (WT), n = 9 GFAP-TK). L) Arrest of cytogenesis significantly worsened recovery of motor function. Time x group interaction F(6, 120) = 3.61, p = 0.0025. *p < 0.05, **p < 0.01, Bonferroni tests. M) Lesion volume was not different between groups. t(20) = 0.27, p = 0.787. N) Lesion reconstruction. Darker shades represent more overlap between animals. Data are presented as mean ± SEM. Where individual datapoints are shown, datapoints representing males are shown as circles; datapoints representing females are shown as squares.

Following GCV administration, we compared motor recovery between GFAP-TK mice and wildtype littermates after photothrombotic cortical infarcts targeting the forelimb area of motor cortex (Tennant et al., 2011; Williamson et al., 2021) (Figure 2K). Motor function was assessed with the single seed reaching task, a highly sensitive and translationally relevant measure of skilled reaching (Klein et al., 2012; van Lieshout et al., 2021). There was no difference between groups in reaching performance during pre-stroke GCV delivery (Bonferroni- corrected p > 0.999). Mice lacking cytogenesis showed significantly worse recovery out to four weeks following stroke (Figure 2L). Lesion size and location did not differ between groups (Figure 2M, N). These results demonstrate that SVZ cytogenesis promotes functional recovery after stroke.

### Aging diminishes the SVZ cytogenic response to stroke and its functional benefits

Aging is associated with a substantially higher incidence of stroke (Kissela et al., 2012) and reduced recovery of function (Paolucci et al., 2003). In animals, aging is associated with diminished SVZ cytogenesis (Bouab et al., 2011; Jin et al., 2004; Kalamakis et al., 2019; Luo et al., 2006). We examined the impact of aging on normal and post-stroke SVZ cytogenesis by comparing young adult (aged 3-6 months) and aged (12-16 months) *Nestin-CreER*; Ai14 mice. Mice were either uninjured (naïve) or subjected to cortical stroke two weeks prior to tissue collection. In young mice, stroke increased the number of Ki67^+^ proliferative cells and tdTomato^+^Sox2^+^ precursors in the SVZ relative to young naïve, aged naïve, and aged stroke mice (Figure 3A-D). By contrast, in aged mice, there was no significant difference in the number of SVZ Ki67^+^ or tdTomato^+^Sox2^+^ cells after stroke relative to naïve mice (p = 0.604, p = 0.998, respectively, Tukey tests). Thus, stroke increases SVZ proliferation and expands the precursor cell pool in young, but not aged, mice.

**Figure 3.**
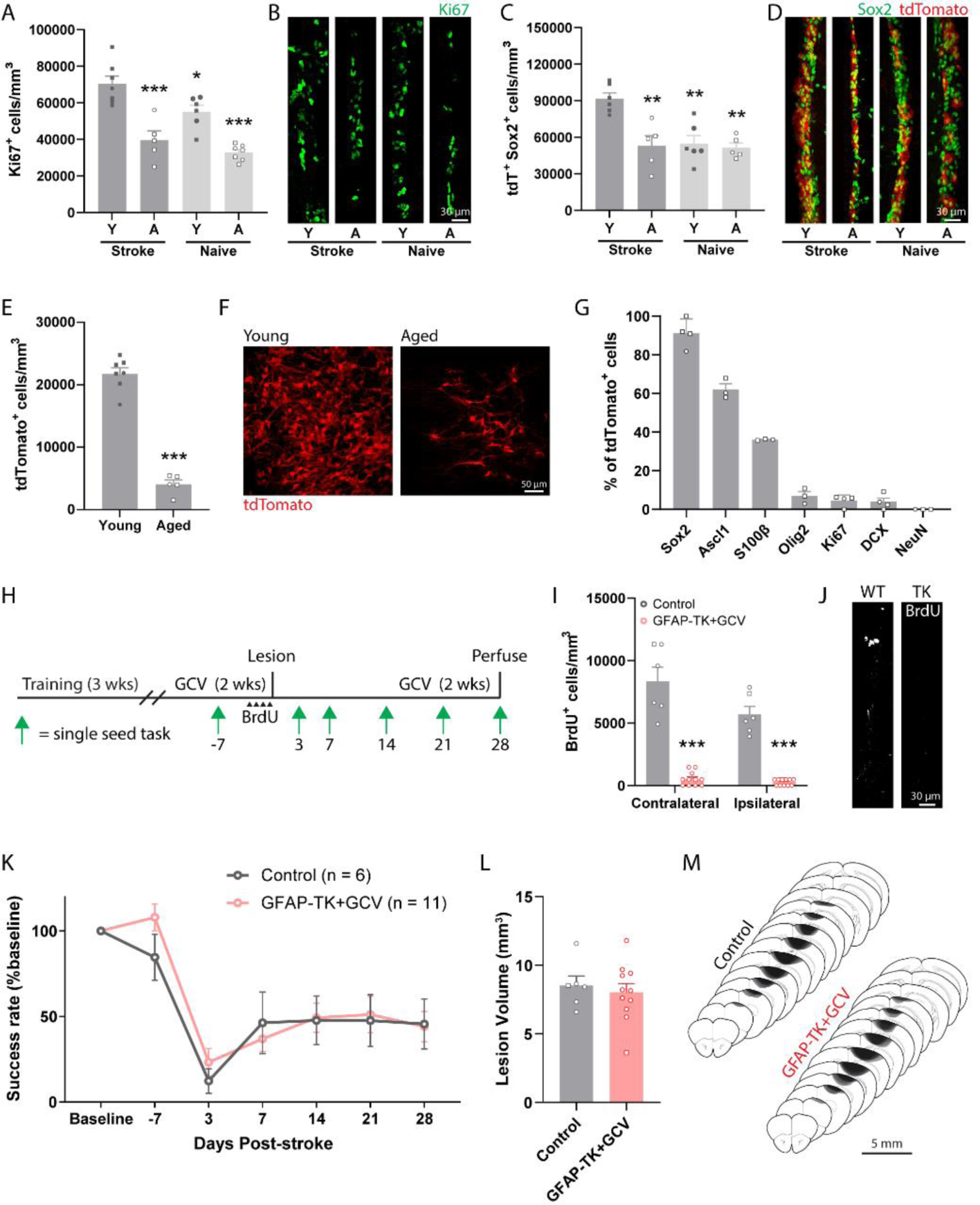
Aging diminishes the cytogenic response to stroke and its contribution to recovery. A-D) Assessment of SVZ cytogenesis in young (Y, 3-6 months) and aged (A, 12-16 months) Nestin-CreER; Ai14 mice (n = 5-7 mice/group). Tissue was collected two weeks post-stroke. The number of proliferating cells (Ki67^+^, A and B) and neural precursor cells (tdTomato^+^ Sox2^+^, C and D) in the SVZ was significantly higher in young mice after stroke compared to all other groups. *p < 0.05, ** p < 0.01, ***p < 0.001 relative to young stroke mice, Tukey tests. E) Fewer tdTomato^+^ cells were observed in peri-infarct cortex of aged mice two weeks post-stroke. ***t(10) = 13.4, p < 0.001. F) Representative images of tdTomato^+^ cells in peri-infarct cortex. G) Quantification of lineage marker expression by tdTomato^+^ cells in peri-infarct cortex of aged mice at two weeks post-stroke (n = 3-4 mice per marker). Similar to what was observed in young mice (Figure 1), most SVZ-derived cells were undifferentiated precursors or astrocytes. H) Experimental timeline for examining the effects of neural stem cell ablation in aged mice (n = 6 controls, n = 11 GFAP-TK given GCV). I) GFAP-TK+GCV mice had fewer BrdU^+^ cells in the SVZ, validating stem cell ablation. ***t(15) ≥ 9.36, p < 0.001. BrdU was given twice per day for two days prior to stroke. J) Representative images of BrdU immunostaining in the SVZ. K) Both aged controls and GFAP-TK+GCV mice showed poor recovery following stroke. There was no difference between groups. Group main effect F(1,15) = 0.17, p = 0.690. L) Lesion volume was not different between groups (t(15) = 0.50, p = 0.628). M) Lesion reconstruction. Darker shades represent more overlap between animals. Data are presented as mean ± SEM. Where individual datapoints are shown, datapoints representing males are shown as circles; datapoints representing females are shown as squares.

We next examined the effects of aging on the migratory response of lineage-traced SVZ cells after stroke. The density of tdTomato^+^ cells in peri-infarct cortex two weeks post-stroke was about five-fold lower in aged mice relative to young mice (Figure 3E, F). Despite the diminished cytogenic response, the phenotype distribution of tdTomato^+^ cells in peri-infarct cortex of aged mice (Figure 3G) was similar to what we observed in young adult mice (Figure 1), with the majority of cells identified as precursors and astrocytes. Thus, far fewer SVZ-derived cells localize to peri-infarct regions in aged mice, but their phenotype distribution is similar to that of young mice.

We next investigated whether the blunted cytogenic response observed in aged mice was still functionally beneficial (Figure 3H). We used GFAP-TK mice to selectively ablate neural stem cells prior to stroke. BrdU pulse labeling of proliferating cells revealed a near complete loss of BrdU^+^ cells in the SVZ of GFAP-TK+GCV mice relative to control aged mice (Figure 3I, J; all mice aged 12-16 months). We assessed motor recovery with the single-seed task and found no differences between groups (Figure 3K). Notably, both aged groups performed significantly worse on day 28 than young mice within intact cytogenesis (data from Figure 2; F(3, 35) = 7.46, p < 0.001 one-way ANOVA; p < 0.012, Tukey tests), but similarly to young GFAP-TK+GCV mice (p > 0.978, Tukey tests). Lesion size was not different between aged groups (Figure 3L, M), and was similar to that of young adults (Figure 2M, N). Altogether, our results indicate that aging diminishes the cytogenic response to stroke and its functional benefits. Together with our findings in young mice, our results show that reduced SVZ cytogenesis, by neural stem cell ablation or aging, is associated with worse functional recovery. Reduced cytogenesis may contribute to worse outcome after stroke with aging.

### Arrest of cytogenesis disrupts neuronal and vascular repair

Our results indicate that most SVZ-derived cells are undifferentiated precursors and reactive astrocytes. We hypothesized that these cell types may influence repair processes to promote behavioral improvement. In particular, synaptic plasticity and vascular remodeling are two functionally important aspects of neural repair that could potentially be augmented by factors produced by SVZ-derived cells. Indeed, past studies have demonstrated beneficial effects of transplanted stem cells and resident cortical astrocytes on these processes (Andres et al., 2011; Bacigaluppi et al., 2016; Horie et al., 2011; Lin et al., 2017; Llorente et al., 2021; Pluchino et al., 2005; Roitbak et al., 2011; Sabelström et al., 2013; Williamson et al., 2021).

We investigated the consequences of neural stem cell ablation on synaptic and vascular plasticity in residual cortex surrounding photothrombotic infarcts with a longitudinal imaging approach (Figure 4A, B). We bred GFAP-TK mice with Thy1-GFP mice, which have sparse, GFP-labeled pyramidal neurons. Resulting double transgenic mice allowed us to monitor synaptic remodeling at single-synapse resolution with repeated 2-photon imaging of dendritic spines on apical dendrites before and after stroke (Brown et al., 2007; Clark et al., 2019; Mostany et al., 2010) with the ability to conditionally arrest cytogenesis. Blood flow was tracked with multi-exposure speckle imaging (MESI), a quantitative, optical, contrast-free technique that yields high-resolution blood flow maps (He et al., 2020; Williamson et al., 2020). Prior to stroke, control and TK^+/-^ mice were unilaterally implanted with cranial windows, trained on the single- seed reaching task, administered GCV or saline, and subjected to baseline imaging. We then induced strokes in forelimb motor cortex and periodically imaged dendritic spines and blood flow and assessed behavioral performance during recovery.

**Figure 4.**
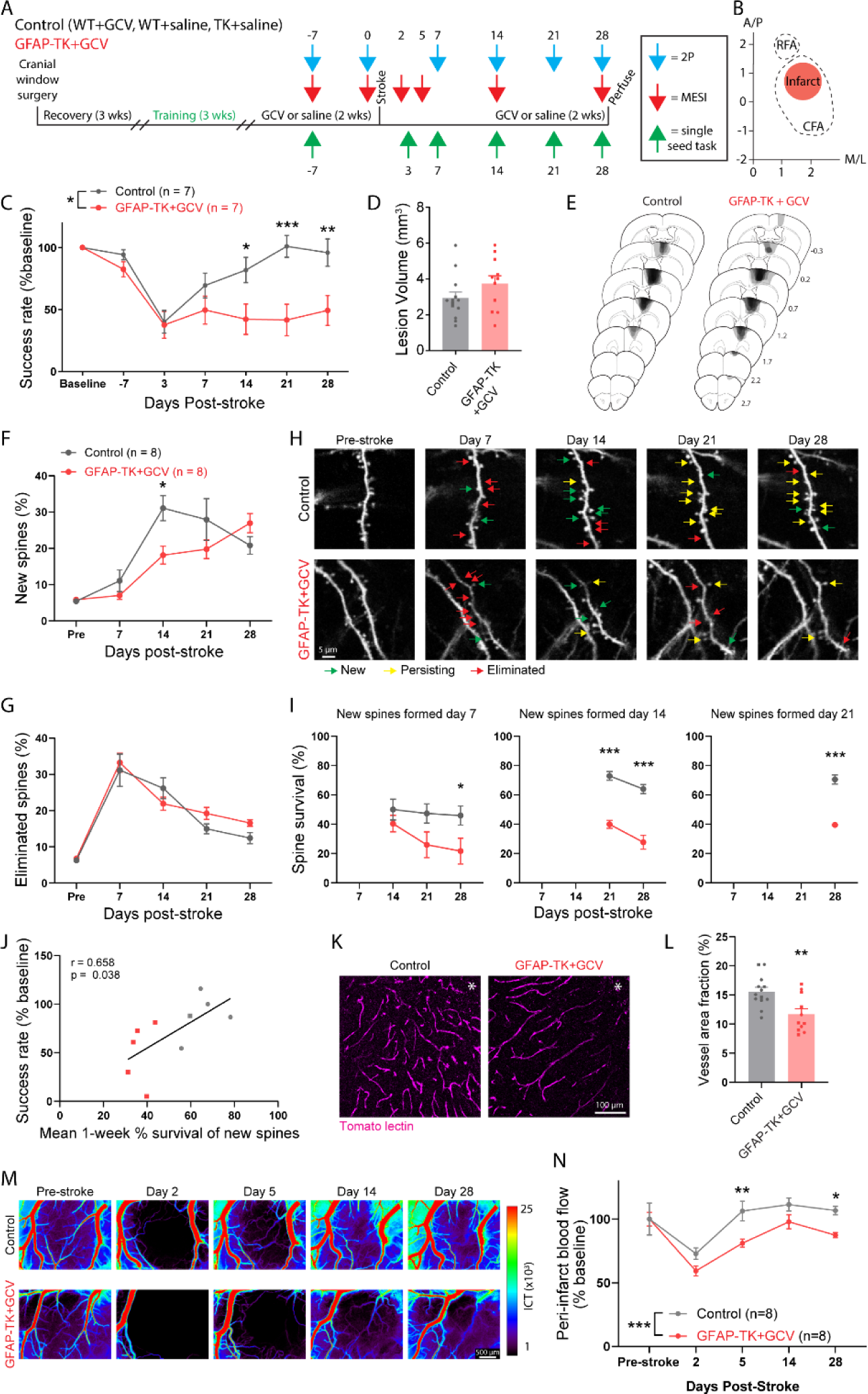
SVZ cytogenesis supports neuronal and vascular remodeling. A) Experimental design for longitudinal imaging and behavioral testing. Subgroups of mice were subjected to longitudinal imaging and behavioral testing (n = 5/group), imaging only (n = 3/group), behavioral testing only (n = 2/group), and neither (histology only; n = 3 control, n = 1 GFAP-TK+GCV). B) Schematic illustrating infarct placement relative to caudal (CFA) and rostral (RFA) forelimb areas in motor cortex. Axes indicate mm relative to Bregma. C) Performance on the single-seed reaching task. GFAP-TK+GCV mice showed significantly worse recovery relative to control mice (significant group x time interaction, F(6, 72) = 4.7, p < 0.001). *p < 0.05, **p < 0.01, ***p < 0.001, Bonferroni tests. D) Lesion volume was not different between groups (t(22) = 1.5, p = 0.159). E) Lesion reconstructions. Darker shades represent more overlap between animals. F-J) Longitudinal imaging of dendritic spines revealed altered spine dynamics in mice with ablated neural stem cells. F) New spine formation was significantly higher in control mice on day 14 (significant time x group interaction, F(4,52) = 4.1, p = 0.006). *p < 0.05, Sidak’s multiple comparison tests between groups for each day. G) Spine elimination was not significantly different between groups (group effect, F(1, 66) = 1.2, p = 0.285). H) Longitudinal 2-photon images of GFP-expressing dendritic spines illustrating spine formation (green arrows), persistence of new spines (yellow arrows), and spine elimination (red arrows). I) Survival of new spines was significantly reduced in GFAP-TK+GCV mice. Plots show survival of new spines formed on days 7, 14, and 21 after stroke. *p < 0.05, ***p < 0.001, Sidak’s multiple comparison tests. J) Spine survival was significantly positively correlated with behavioral performance on the final day of testing. K) Representative images of peri-infarct vasculature. L) Peri-infarct vessel density was significantly reduced in GFAP-TK+GCV mice (**t(22) = 3.3, p = 0.004). M) Representative MESI images of blood flow. N) Peri-infarct blood flow was significantly reduced on days 5 and 28 in GFAP-TK+GCV mice (group effect, F(1, 14) = 32.98, p < 0.001). *p < 0.05, **p < 0.01, Sidak’s multiple comparison tests. Data are presented as mean ± SEM. Where individual datapoints are shown, datapoints representing males are shown as circles; datapoints representing females are shown as squares.

Motor recovery was again significantly impaired in GFAP-TK+GCV mice, with no differences in lesion size observed between groups (Figure 4C-E). 2-photon imaging of dendritic spines revealed increased spine turnover in peri-infarct cortex (Figure 4F-H), consistent with past work (Brown et al., 2007; Clark et al., 2019; Joy et al., 2019; Mostany et al., 2010). There were no group differences in spine dynamics before stroke (Supplemental Figure 4). New spine formation peaked during the second week after stroke, and was significantly higher in mice with intact cytogenesis. Spine elimination was greatest during the first week post-stroke, and subsequently declined with time, without significant differences between groups. A subpopulation of new spines formed after stroke persists long-term, and the persistence of newly formed spines is associated with greater functional recovery (Clark et al., 2019). The survival of new spines was significantly reduced in GFAP-TK+GCV mice regardless of the day of spine formation (Figure 4I). Moreover, spine survival rate was positively correlated with behavioral performance on the final assessment day (Figure 4J). Thus, SVZ cytogenesis supports synaptic remodeling after stroke, particularly by promoting the long-term stabilization of new synapses.

Broad regions of reduced blood flow persist for days to weeks surrounding focal strokes (He et al., 2020; Williamson et al., 2020). Remodeling of peri-infarct vasculature helps to restore blood flow and is associated with behavioral improvement (Williamson et al., 2020). To examine vascular remodeling, we injected mice with fluorophore-conjugated tomato lectin immediately before euthanasia to label perfused vasculature. Vessel density in peri-infarct cortex was significantly reduced in GFAP-TK+GCV mice relative to controls (Figure 4K, L). Furthermore, longitudinal blood flow imaging demonstrated impaired recovery of blood flow at days 5 and 28 post-stroke (Figure 4M, N). Early vascular permeability was not affected by arresting cytogenesis (Supplemental Figure 3). Overall, these findings demonstrate that SVZ-derived cells beneficially shape neuronal and vascular remodeling processes after stroke in order to promote recovery.

### SVZ-derived cells interact with vasculature and produce trophic factors

Neuroblasts originating in the SVZ migrate along vascular scaffolds towards the olfactory bulb in the healthy brain (Bovetti et al., 2007) and towards peri-infarct regions after stroke (Ohab et al., 2006; Thored et al., 2007). We observed frequent contact between lineage- traced SVZ progeny and blood vessels in peri-infarct cortex, including contact of nearby vessels by the processes or cell bodies of 88.2% of SVZ-derived cells (Figure 5A-C). Thus, contact with blood vessels may underlie some of the reparative effects of SVZ-derived cells. In addition, stroke may induce expression of migratory cues in endothelial cells to drive migration of reparative cells from the SVZ (Ohab et al., 2006).

**Figure 5.**
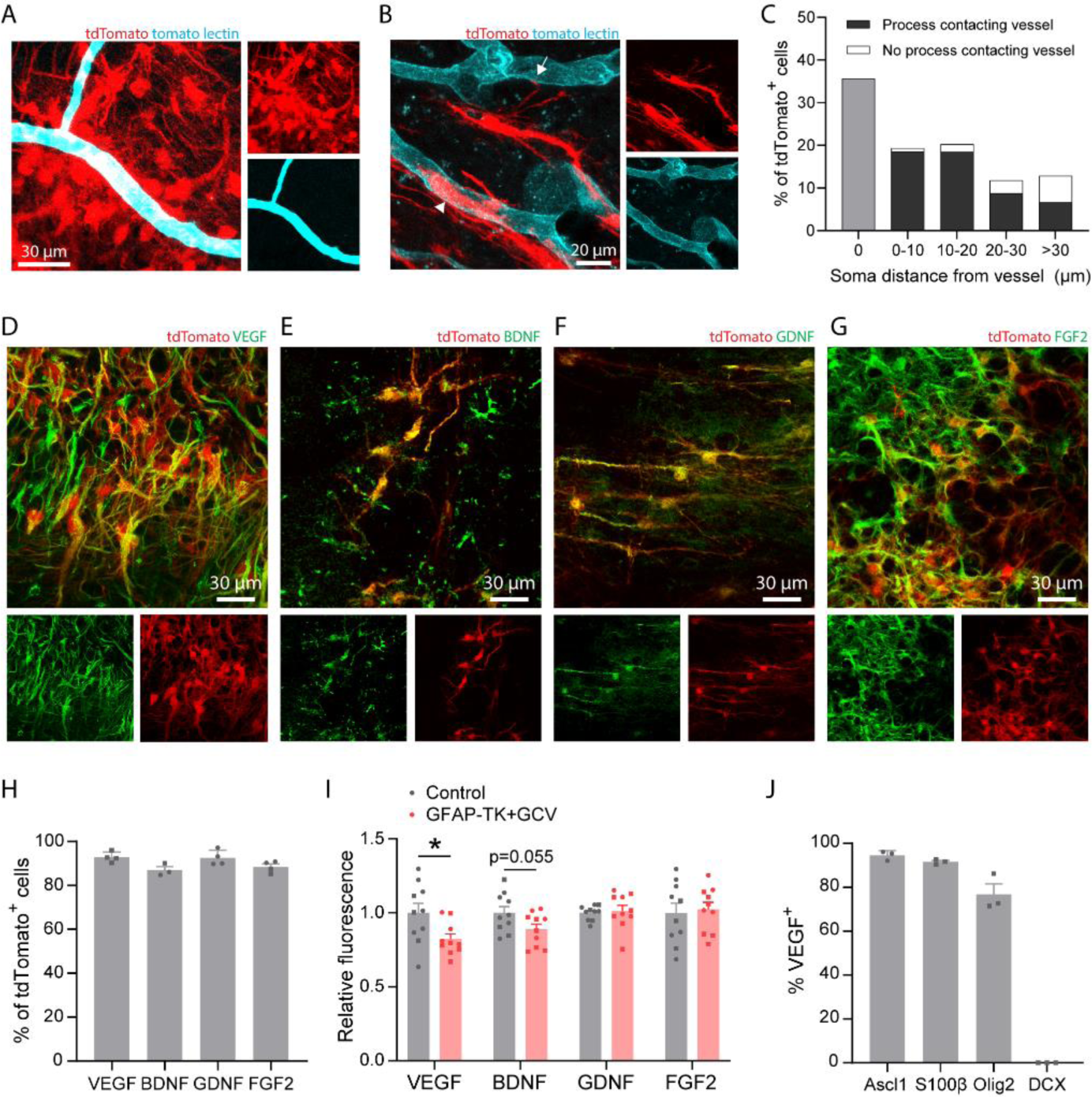
SVZ-derived cells interact with vasculature and produce trophic factors. A) Confocal image showing clustering of lineage traced cells around blood vessels in peri-infarct cortex. B) tdTomato-expressing cells were frequently observed with cell bodies abutting vessels (arrowhead) or extending processes that terminated on nearby vessels (arrow). C) Quantification of SVZ-derived cell interaction with vasculature in peri-infarct cortex two weeks post-stroke. Data are from 661 cells across 3 Nestin-CreER; Ai14 mice. D-G) Confocal images from peri-infarct cortex showing expression of the trophic factors VEGF (D), BDNF (E), GDNF (F), and FGF2 (G) in tdTomato^+^ cells. H) Quantification of trophic factor expression in tdTomato^+^ cells. I) Quantification of trophic factor expression in peri-infarct cortex four weeks post-stroke between control mice and GFAP-TK+GCV mice (n = 10/group). Fluorescence intensity is reported relative to controls. VEGF protein fluorescence was significantly reduced in GFAP- TK+GCV mice. *t(18) = 2.4, p = 0.025. J) Quantification of VEGF expression by phenotype in lineage traced cells. VEGF was expressed by Ascl1^+^ precursors, S100β^+^ astrocytes, and Olig2^+^ oligodendrocyte-lineage cells, but not by neuronal lineage cells (DCX^+^). Data are presented as mean ± SEM. Where individual datapoints are shown, datapoints representing males are shown as circles; datapoints representing females are shown as squares.

Transplanted neural precursors of various sources and in diverse disease settings have been reported to express trophic factors, which may be implicated in their therapeutic effects (Andres et al., 2011; Bacigaluppi et al., 2009; Drago et al., 2013; Horie et al., 2011; Llorente et al., 2021; Martino and Pluchino, 2006; Roitbak et al., 2011). To prospectively investigate molecular mechanisms underlying the facilitation of post-stroke repair by SVZ cytogenesis, we examined the expression of four trophic factors known to drive neuronal and vascular growth.

∼90% of all SVZ-derived lineage traced cells in peri-infarct cortex expressed VEGF, BDNF, GDNF, and FGF2 (Figure 5D-H). We compared relative abundance of each of these proteins in peri-infarct cortex between control and GFAP-TK+GCV mice at 28 days post-stroke to evaluate production of these factors by SVZ-derived cells relative to other cell types. VEGF protein was significantly reduced in GFAP-TK+GCV mice, indicating that SVZ-derived cells are a major source of VEGF (Figure 5I). Notably, among SVZ-derived cells, all lineages except for new neurons produced VEGF (Figure 5J). These findings suggest that SVZ cytogenesis may facilitate neural repair at least in part through production of trophic factors, particularly VEGF.

### Conditional deletion of Vegf in adult neural stem cells impairs recovery and repair

VEGF promotes the growth of blood vessels and neurons (Gerber et al., 1999; Raab et al., 2004; Rosenstein et al., 2003; Sun et al., 2003). We next investigated whether VEGF produced by SVZ-derived cells was involved in post-stroke recovery and repair. We generated *Nestin-CreER*; *Vegf*^fl/fl^ mice to permit inducible deletion of *Vegf* in adult neural stem cells and their progeny (VEGF cKO; Figure 6A). Lineage tracing in *Nestin-CreER*; *Vegf*^fl/fl^; Ai14 mice showed a near complete loss of VEGF in tdTomato^+^ cells in peri-infarct cortex, but no change in the number of migratory cells relative to controls at two weeks post-stroke (Figure 6B-E).

**Figure 6.**
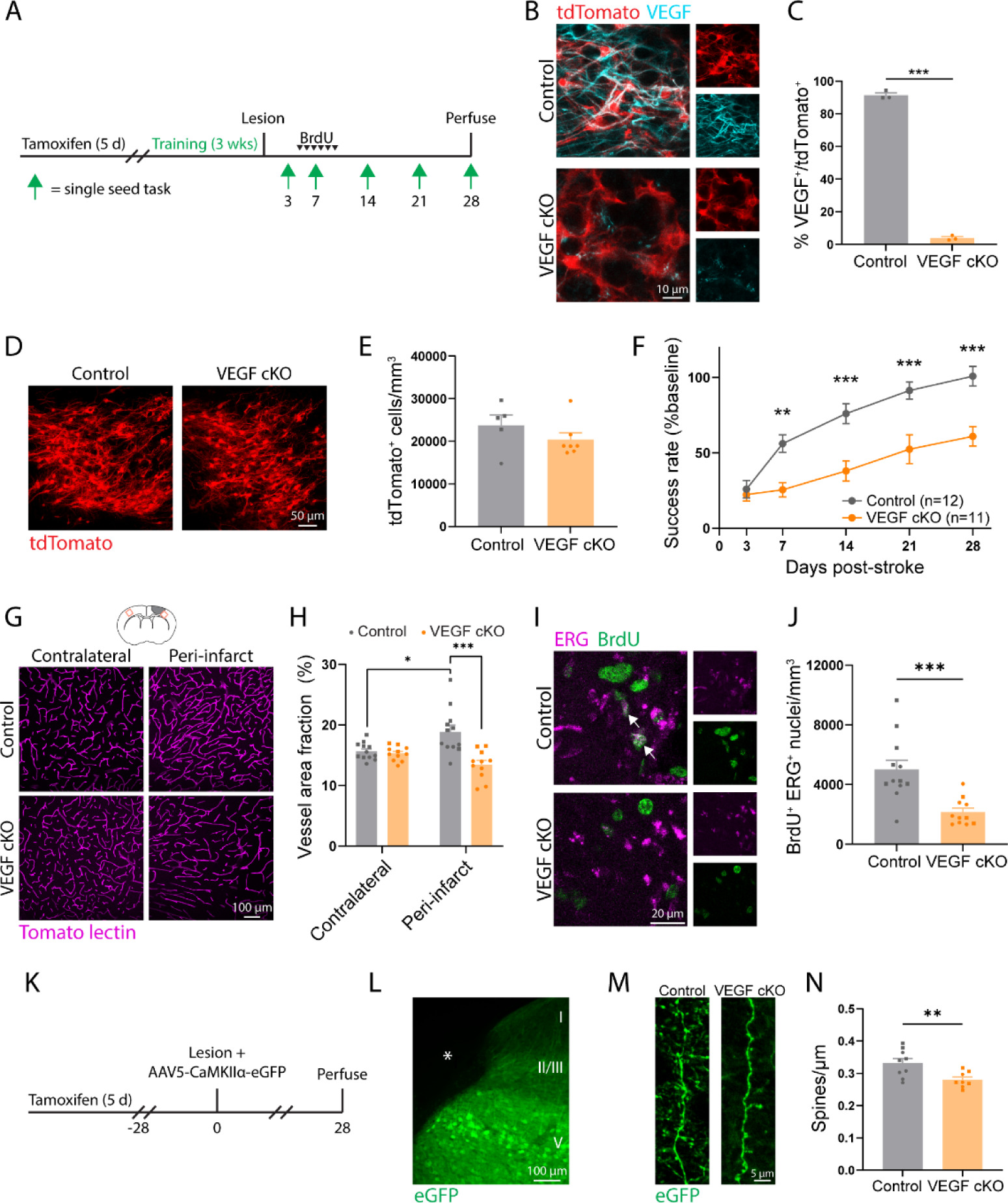
Adult neural stem cell-specific deletion of *Vegf* impedes recovery and repair. A) Timeline of experiments to examine recovery in control (Nestin-CreER^+/-^; VEGF^fl/+^, Nestin- CreER^-/-^; VEGF^fl/+^, or Nestin-CreER^+/-^; VEGF^+/+^) and VEGF cKO (Nestin-CreER^+/-^; VEGF^fl/fl^) mice. B) Confocal images demonstrating VEGF loss in tdTomato^+^ cells in Nestin-CreER^+/-^; VEGF^fl/fl^; Ai14 mice. Images are from peri-infarct cortex two weeks post-stroke. C) Substantially fewer tdTomato^+^ cells express VEGF in VEGF cKO mice. n = 3 mice/group. ***t(4) = 51.23, p < 0.001. D) Representative images of tdTomato^+^ cells in peri-infarct cortex two weeks post-stroke. (E) There was no difference in peri-infarct tdTomato^+^ cell density between controls and VEGF cKO mice, indicating that the cytogenic response was unaffected. N = 5 controls, n = 7 VEGF cKO. t(10) = 1.18, p = 0.266. F) VEGF cKO significantly impaired motor recovery measured with the single-seed reaching task (n= 12 control, n = 11 VEGF cKO). Time x group interaction F(4, 84) = 9.2, p < 0.001. **p < 0.01, ***p < 0.001, Sidak’s tests between group for each timepoint. G) Representative confocal images and quantification (H) of vasculature show that VEGF cKO markedly reduced peri-infarct vessel density relative to control mice. ***t(21) = 4.1, p < 0.001. Within control mice, peri-infarct vessel density was significantly greater than in contralateral cortex, consistent with stroke-induced neovascularization. *t(22) = 2.8, p = 0.011. By contrast, VEGF cKO mice had diminished vessel density in peri-infarct cortex relative to the intact contralateral cortex, indicating a failure of neovascularization (t(20) = 2.3, p = 0.030). I) Representative confocal images of new endothelial cells (BrdU^+^ ERG^+^) in peri-infarct cortex. J) The number of BrdU^+^ ERG^+^ nuclei was significantly less in VEGF cKO mice. ***t(15.0) = 4.2, p < 0.001, Welch’s corrected t-test. K) Timeline of experiments for evaluating peri-infarct spine density in control (n =9) and VEGF cKO (n = 8) mice. L) Layer V pyramidal neurons were labeled by intracortical injections of AAV5-CaMKiiα-eGFP. Image shows eGFP labeling in peri-infarct cortex. Asterisk indicates lesion core. M) Representative images and quantification (N) of dendritic spine density. Spine density was significantly lower in VEGF cKO mice. **t(15) = 3.0, p = 0.009. Apical dendrites were sampled from layer II/III between 100-700 μm from the infarct border. 2631 spines were counted along 7.9 mm total length of dendrite in controls. 1934 spines were counted along 6.9 mm total length of dendrite in cKO mice. Data are presented as mean ± SEM. Where individual datapoints are shown, datapoints representing males are shown as circles; datapoints representing females are shown as squares.

Recovery of forelimb motor function was significantly worse in VEGF cKO mice relative to controls as measured with the single-seed reaching task (Figure 6F). Lesion size and location were not different between groups (Supplemental Figure 5). Vascular remodeling following stroke is seen by increases in peri-infarct vascular density (Williamson et al., 2021, 2020). While vessel density in peri-infarct cortex was increased in control mice relative to the contralateral hemisphere, this was not seen in VEGF cKO mice, and peri-infarct vessel density in VEGF cKO mice was reduced relative to controls (Figure 6G, H). We injected mice with BrdU daily during the peak in angiogenesis from days 5-10 after stroke to quantify new vessel formation. The number of new endothelial cells (BrdU^+^ERG^+^) was significantly reduced in VEGF cKO mice, confirming impaired angiogenesis in mice lacking VEGF in SVZ-derived cells.

In a separate experiment, we examined the effects of conditional *Vegf* deletion on dendritic spine density after stroke (Figure 6K). Peri-infarct layer V cortical pyramidal neurons were labeled by injections of AAV5-CaMKIIa-eGFP (Figure 6L). Four weeks post-stroke, we quantified spine density on apical dendrites in layer II/III. We previously found that changes in spine density in this region parallel functional recovery (Clark et al., 2019). Spine density was significantly reduced in VEGF cKO mice (Figure 6M, N). Altogether, our results identify VEGF produced by SVZ-derived cells as a key driver of repair and recovery after stroke. More broadly, our findings position newborn cells arising from the SVZ in response to injury as a unique cellular source of trophic support that instructs neural repair.

### AAV-mediated expression of VEGF in peri-infarct cortex enhances recovery in mice with arrested cytogenesis

We next tested whether replacement of VEGF would be sufficient to enhance recovery in mice in which cytogenesis was arrested. GFAP-TK mice were trained on the single-seed reaching task and administered GCV to ablate neural stem cells. Immediately after stroke, mice were injected with either AAV5-EF1α-VEGF-P2A-eGFP (AAV-VEGF-eGFP), to induce VEGF and eGFP expression, or AAV5-EF1α-eGFP (AAV-eGFP), to induce only eGFP expression, into layer V of peri-infarct cortex (Figure 7A-C). AAV-VEGF-eGFP induced rapid and sustained motor recovery as measured by the single-seed reaching task, whereas AAV-eGFP injected mice showed little improvement up to four weeks post-stroke (Figure 7D). Lesion size and location were not different between groups (Supplemental Figure 6). We examined vascular density in homotopic contralateral and peri-infarct cortex 28 days post-stroke. Vascular density was significantly greater in peri-infarct cortex of AAV-VEGF-eGFP injected mice relative to AAV-eGFP mice (Figure 7E, F). Moreover, there was no difference in vascular density between contralateral and peri-infarct cortex in the AAV-eGFP group, indicating a failure of neovascularization (t(22) = 0.97, p = 0.344). Mice were given daily injections of BrdU during days 5-10 post-stroke to label new blood vessels. AAV-VEGF-eGFP mice had substantially more angiogenesis in peri-infarct cortex than AAV-eGFP mice as measured by the number of BrdU^+^ERG^+^ nuclei (Figure 7G, H). Finally, we examined spine density of eGFP-expressing pyramidal neurons on apical dendrites in layer II/III. Spine density was significantly higher in mice given AAV-VEGF-eGFP (Figure 7I, J). In wildtype mice subjected to a sham stroke procedure, AAV-VEGF-eGFP increased vessel density but did not affect motor function (Supplemental Figure 7). Overall, our findings indicate that replacement of VEGF is sufficient to enhance repair and recovery in mice lacking cytogenesis. More broadly, these findings suggest that replacement of factors produced by SVZ-derived cells may constitute an effective therapy.

**Figure 7.**
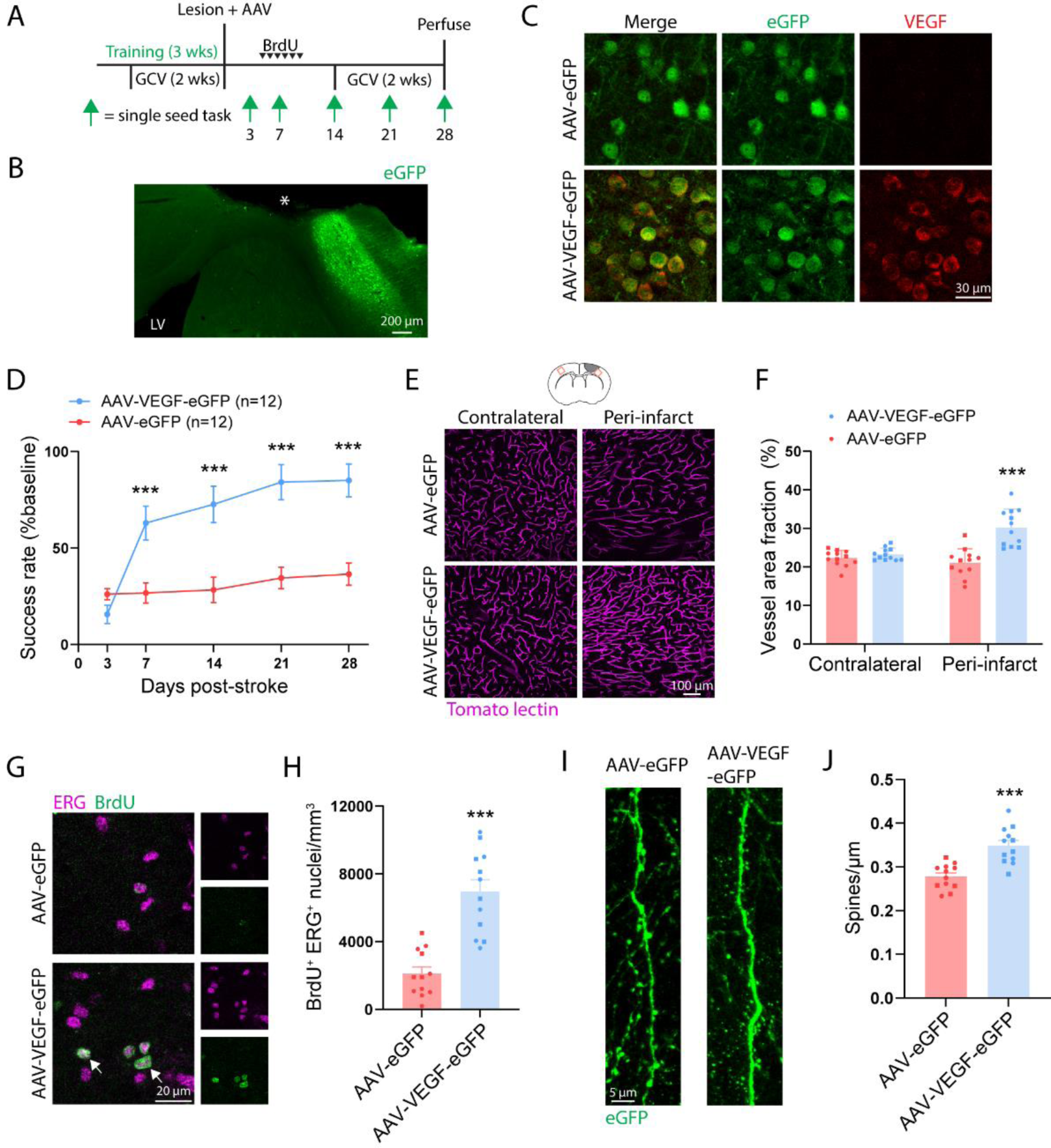
VEGF rescues poor recovery due to neural stem cell ablation. A) Experimental timeline. Both groups used GFAP-TK mice given GCV to ablate neural stem cells (n = 12 mice per group). Animals were injected with either AAV5-Ef1α-eGFP (AAV-eGFP) or AAV5-Ef1α-VEGF-P2A-eGFP (AAV-VEGF-eGFP) in layer V of peri-infarct cortex. B) Image illustrating AAV targeting of peri-infarct cortex. Asterisk indicates lesion core. LV, lateral ventricle. C) Confocal images validating that AAV-VEGF-eGFP induces VEGF expression. D) AAV-VEGF-eGFP improved motor recovery on the single seed reaching task. Significant time x group interaction F(4, 88) = 15.9, p < 0.001. **p < 0.01, ***p < 0.001, Sidak’s multiple comparison tests. E) Representative confocal images and quantification (F) of vasculature show that AAV-VEGF-eGFP increased peri-infarct vessel density relative to AAV-eGFP mice. ***t(22) = 5.3, p < 0.001, t test. G) Representative confocal images of new endothelial cells (BrdU^+^ ERG^+^, arrows) in peri-infarct cortex. H) The number of BrdU^+^ ERG^+^ nuclei was significantly greater in AAV-VEGF-eGFP mice. ***t(22) = 6.0, p < 0.001. I) Representative images and quantification (J) of dendritic spine density. Spine density was significantly higher in AAV-VEGF-eGFP mice. ***t(22) = 4.8, p < 0.001. Apical dendrites were sampled from layer II/III between 100-800 μm from the infarct border. 2825 spines were counted along 10.2 mm total length of dendrite in AAV-eGFP mice. 3231 spines were counted along 9.3 mm total length of dendrite in AAV-VEGF-eGFP mice. Data are presented as mean ± SEM. Where individual datapoints are shown, datapoints representing males are shown as circles; datapoints representing females are shown as squares.

## Discussion

Our study revealed that a previously underappreciated class of cells, undifferentiated precursors, constitutes the majority of cells that arise from the SVZ and migrate towards the site of injury following stroke. The migration of primarily undifferentiated cells towards the site of injury suggests that the main function of post-injury cytogenesis is likely not cell replacement. We found that reducing SVZ cytogenesis, by neural stem cell ablation or aging, leads to poor functional recovery. Moreover, synaptic and vascular repair were disrupted in mice with deficient cytogenesis. SVZ-derived cells produced trophic factors, most notably VEGF. Loss-of-function experiments demonstrated that VEGF produced by SVZ-derived cells is crucial for effective repair and recovery. Finally, gain-of-function experiments showed that replacement of VEGF was sufficient to enhance recovery in mice lacking cytogenesis. We conclude that trophic support from SVZ-derived cells drives neural repair and functional recovery after stroke. Thus, newborn cells formed in response to injury enable recovery by acting as a unique source of trophic cues that instruct neural repair.

With lineage tracing of adult neural stem cells and extensive phenotyping, we identified undifferentiated precursors as the largest subpopulation of SVZ-derived cells after stroke.

Undifferentiated neural precursor cells may the better suited than differentiated neural cell types to facilitate neural repair. In culture, neural precursors secrete factors that facilitate vessel formation and neuronal outgrowth (Kirby et al., 2015; Roitbak et al., 2011; Rosenstein et al., 2003). In addition, transplantation of neural stem cells enhances repair and recovery in models of stroke without differentiation of transplanted cells (Andres et al., 2011; Bacigaluppi et al., 2016; Horie et al., 2011; Llorente et al., 2021; Roitbak et al., 2011). Collectively, these studies illustrate the reparative abilities of undifferentiated precursors and suggest that post-stroke cytogenesis facilitates neural repair without substantial cell replacement.

Further work is needed to understand how the fate of individual cells is decided among the total population of newborn cells arising from the SVZ. Principles of neural development may apply to cell fate decisions during post-injury cytogenesis in adulthood. For example, the transcription factor NFIA controls gliogenesis during development (Deneen et al., 2006), and is also necessary for SVZ astrogenesis after stroke (Laug et al., 2019). In addition, NFIA inhibits neurogenesis via Notch effectors (David-Bercholz et al., 2021; Deneen et al., 2006).

Accordingly, interfering with Notch signalling in neural stem cells biases SVZ progeny towards a neuronal fate and away from an astrocytic fate after stroke (Benner et al., 2013). Additional work will be needed to clarify the mechanisms that dictate phenotypes of SVZ cells responding to stroke, and whether different cell types have distinct reparative functions.

Our finding that aging reduces SVZ cytogenesis could be translationally relevant given that stroke incidence increases with age (Kissela et al., 2012). We observed the loss of stroke-induced SVZ proliferation and precursor cell pool expansion in aged mice, which suggests deficient activation of neural precursor cells with aging. Inflammatory signals have been implicated in controlling quiescence/activation of precursors. Chronic inflammatory signals, including interferons, promote stem cell quiescence in aging (Kalamakis et al., 2019). By contrast, acute interferon signaling after stroke stimulates precursor activation (Belenguer et al., 2021; Llorens-Bobadilla et al., 2015). Therefore, inflammatory signaling pathways may be a target to restore cytogenesis in aged animals. We also observed a reduction in the number of cells localized in peri-infarct regions in aged mice that was disproportionate relative to the diminishment of SVZ cytogenesis. Thus, there may also be an age-dependent reduction in either the migratory ability of SVZ cells or the expression of migratory cues at the site of injury.

SVZ-derived cells interacted closely with peri-infarct blood vessels and produced trophic factors to facilitate their growth after stroke. The ectopic migration of cells from the SVZ is caused in part by expression of migratory cues in peri-infarct vasculature after stroke (Ohab et al., 2006; Thored et al., 2007). This may be an adaptive response to attract precursors that facilitate tissue growth and repair. Similar processes are mirrored in at least two cases during development. First, neuroepithelial cells produce VEGF to stimulate initial embryonic brain angiogenesis (Breier et al., 1992; Raab et al., 2004). Second, endothelial cues drive oligodendrocyte precursor cell attachment and migration along vessels in the embryonic nervous system (Tsai et al., 2016), and oligodendrocyte precursor-derived HIF-dependent factors, including VEGF, subsequently drive early postnatal angiogenesis (Yuen et al., 2014; Zhang et al., 2020). The ectopic migration of reparative SVZ cells towards injury may represent a reuse of tissue growth mechanisms from development.

We have demonstrated that the SVZ cytogenic response to stroke primarily produces undifferentiated precursors that localize to peri-infarct regions – the site of neural repair. SVZ-derived cells produce VEGF that is critical for effective vascular and synaptic plasticity, and ultimately behavioral recovery. Thus, our findings position SVZ cytogenesis as a mechanism that promotes recovery via trophic support rather than cell replacement. These findings provide insight into a fundamental brain repair process and may be relevant for informing treatment strategies.

## Materials and Methods

### Subjects and experimental design

Young adult (3-6 months) and aged (12-16 months) mice of both sexes were used. All mice were on a predominantly C57BL/6 background. Transgenic strains were Rosa-CAG-LSL-tdTomato (Ai14, JAX #007914), Nestin-CreER (JAX #016261), ASCL1-CreER (JAX #012882), Rosa-CAG-LSL-Sun1-sfGFP (JAX #021039), GFAP-TK (JAX #005698), Thy1-GFP M-line (JAX #007788), floxed Vegfa (Genentech) (Gerber et al., 1999) (Supplemental Table 1). Mice were bred locally. Animals were housed 2-5 per cage with free access to food and water, except during periods of restricted feeding for behavioral training and assessment. Animals were randomized to groups except when group assignment was dependent on genotype.

Experimentation and analysis were done blinded to group allocation. Experiments consisted of 1- 5 cohorts of animals. Sample sizes were based on past work using similar methods (Benner et al., 2013; Brown et al., 2007; Clark et al., 2019; Williamson et al., 2021, 2020).

### Drug administration

100 mg/kg of 20 mg/mL tamoxifen dissolved in corn oil was given (i.p.) daily for 5 consecutive days. 100 mg/kg of 10 mg/mL BrdU dissolved in saline was given (i.p.) once or twice per day for 2 or 6 consecutive days. Ganciclovir dissolved in saline was delivered continuously for 14 days via subcutaneous osmotic pumps (Azlet) at a rate of 6.25 µg/hr. In some experiments, a second course of ganciclovir was given beginning two weeks after stroke to maintain stem cell ablation.

### Cranial window implantation

Chronic glass cranial windows were placed over forelimb motor cortex (Clark et al., 2019; Williamson et al., 2021, 2020). Isoflurane (3% induction, 1-2% maintenance) in oxygen was used for anesthesia. Circular craniotomies (∼4.5 mm diameter) were made 1.5 mm lateral from Bregma. 4 mm glass windows (Warner Instruments) were secured in place with cyanoacrylate, and exposed skull was covered with dental cement. Carprofen (5 mg/kg, i.p.) was given daily for 7 days to minimize inflammation.

### Ischemic stroke

To model stroke, unilateral photothrombotic lesions were induced in the forelimb region of motor cortex (Tennant et al., 2011; Williamson et al., 2021). Isoflurane (3% induction, 1-2% maintenance) in oxygen was used for anesthesia. Body temperature was maintained with a heated pad for the duration of anesthesia. Stroke was induced through the intact skill by making a scalp incision, administering rose bengal (0.15 mL, 15 mg/mL, i.p.), and illuminating the skull 2 mm lateral from Bregma with a surgical lamp (Schott KL 200) for 15 minutes though a 3 mm aperture. For animals with cranial windows, penetrating arterioles supplying motor cortex were identified by live speckle contrast imaging, and subsequently targeted with a 20 mW 532 nm laser for 15 minutes after administering rose Bengal (0.2 mL, 15 mg/mL, i.p.) (Williamson et al., 2020). Sham stroke procedures involved omitting either illumination or rose bengal.

### Virus injections

Cortical layer V was targeted for virus injections with a Drummond Nanoject II microinjector through a pulled pipette. Injections were made 0.7 mm below the pial surface at three locations relative to Bregma: 2 mm anterior, 2 mm lateral; 0.5 mm anterior, 3.2 mm lateral; and 1 mm posterior, 2.8 mm lateral. 230 nL was injected per site at a rate of 46 nL/min. The pipette was left in place for 2 minutes after the final injection at each site before it was slowly removed. See Supplemental Table 1 for details on viruses.

### 2-photon imaging of dendritic spines

Mice were anesthetized with isoflurane (3% induction, ∼1.5% maintenance) in oxygen and head-fixed to minimize breathing artifacts. Imaging was done with a Prairie Ultima 2-photon microscope with a Ti:Sapphire laser (MaiTai, Spectra Physics) tuned to 870 nm. Image stacks were acquired with 512 x 512 pixel resolution and 0.7 µm z step size using a water-immersion 20×/1.0 (Olympus) objective. 4x magnification yielded a 117.2 µm x 117.2 µm field of view.

Imaging was done weekly, including two pre-stroke and four post-stroke imaging sessions. During pre-stroke imaging, typically 6-10 regions of unobstructed dendrites were imaged to a depth of ∼150 µm. Regions were selected based on a predicted proximity of <1 mm from the infarct. After stroke, the infarct border was identified from blood flow maps and loss of GFP fluorescence (Supplemental Figure 4). Regions within 700 µm of the infarct border were re-imaged at subsequent time points in order to track spine dynamics during recovery. In some animals, additional imaging regions were added after stroke. Time lapse images of dendrites >30 µm in length with clearly visible spines were analyzed to quantify spine formation and elimination, and persistence of newly formed spines (Clark et al., 2019; Joy et al., 2019; Tennant et al., 2017).

### Blood flow imaging

Blood flow was imaged through cranial windows with multi-exposure speckle imaging, a label-free, quantitative, optical method (Clark et al., 2019; Williamson et al., 2021, 2020).

Anesthetic level was consistent across all imaging sessions (1.25% isoflurane in oxygen). Two pre-stroke images were collected to establish baseline blood flow. Post-stroke images were collected on days 2, 5, 14, and 28. Each imaging session lasted <10 mins. Blood flow was measured in parenchymal regions and tracked over time as before (Williamson et al., 2021).

### Behavioral testing

Skilled forelimb use was assessed with the single seed reaching task, which is highly sensitive to deficits caused by motor cortical damage and is translationally relevant (Clark et al., 2019; Klein et al., 2012; van Lieshout et al., 2021; Williamson et al., 2020). Animals were food restricted to ∼90% free feeding weight to encourage reaching. First, animals were shaped on the task and the preferred paw for reaching was determined over 2-5 days. Training was done over 15 sessions, once per day, five days per week. Each session consisted of 30 trials. For each trial, animals were allowed up to two reach attempts. A successful reach was defined as the animal grasping the seed and bringing it to its mouth. Failure was defined as missing the seed, knocking it out of the well, or releasing it before it was brought to the animal’s mouth. Baseline performance was defined as the mean success rate per trial over the last three training sessions. Inclusion criteria was a minimum baseline success rate of 30%. 1 mouse (GFAP-TK+GCV) from the experiment in Figure 2,8 mice (n = 7 control, n = 1 GFAP-TK+GCV) from the experiment in Figure 3, and 2 mice (n = 1 AAV-eGFP, n = 1 AAV-VEGF-eGFP) from the experiment in Supplemental Figure 7 failed to meet this threshold and were excluded. Test sessions were done on days 3, 7, 14, 21, and 28 post-stroke.

### Histology and image analysis

Mice were euthanized by overdosed with a pentobarbitol/phenytoin solution followed by perfusion with 0.1M phosphate buffer and 4% paraformaldehyde in phosphate buffer. Brains were postfixed overnight at 4°C. 35 µm coronal sections were collected with a vibratome (VT1000S, Leica). To label vasculature, mice were retro-orbitally injected with 0.1 mL of Dylight 594- or 649-conjuagated tomato lectin 5 minutes prior to perfusion (Williamson et al., 2021, 2020). To examine vascular permeability, mice were retro-orbitally injected with 0.1 mL of 50 mg/mL FITC-conjugated albumin 2 hours prior to perfusion.

To create lesion reconstructions and quantify lesion volume, one set of every fifth section was Nissl stained. Lesions were reconstructed as previously described (Kim et al., 2018). Lesion volume was calculated using Cavalieri’s method as the difference in volume between uninjured and injured cortex.

Immunohistochemical staining was done by washing tissue in phosphate buffered saline (PBS), blocking with 10% donkey serum in PBS with 0.25% Triton for 60 minutes, incubating with primary antibodies overnight (antibodies and dilutions are reported in Supplemental Table 1), washing in PBS, incubating with species appropriate 405-, 488-, 594, or 647-conjugated secondary antibodies, and washing a final time in PBS. Tissue was pretreated with 2 N HCl (30 mins) followed by 0.1 M boric acid (10 mins) when staining for BrdU.

Confocal images with 1-2 µm Z-step size were collected with a Leica TCS SP5 microscope. 20×/0.7 NA and 40×/1.0NA objectives were used. Acquisition settings were consistent between samples. Typically, 3 sections were imaged per region of interest per mouse.

FIJI was used for image analysis. Area fraction and fluorescence intensity were quantified as before (Williamson et al., 2021). The optical disector method was used to count cell density. Cell density was calculated by number of cells / (frame area × section thickness).

### Statistics

Data are expressed as mean ± S.E.M. Measurements from individual animals are shown on plots as datapoints where possible. Data were analyzed with GraphPad Prism version 9.3. Independent samples were compared with two-tailed t tests. Variance was assessed with F tests, and Welch’s corrected t tests were used in cases where variance was significantly different. One- and two-way ANOVAs, mixed-effects analyses, and linear regressions were used as noted in the text. Post hoc tests were used following significant ANOVA as noted in the text. Details on the statistical tests used for each experiment are located in the Results and figure legends. Alpha was set at P < 0.05.

### Study approval

Animal use was in accordance with a protocol approved by the Institutional Animal Care and Use Committee at the University of Texas at Austin.

## Author contributions

Conceptualization: M.R.W.; Methodology: M.R.W., T.A.J., and M.R.D.; Investigation and analysis: M.R.W., S.P.L., R.L.F., N.A.D., and J.L.R.; Writing – Original Draft: M.R.W.; Writing – Review & Editing: all authors; Supervision: T.A.J. and M.R.D.; Funding Acquisition: A.K.D., M.R.D., and T.A.J.

## Acknowledgments

This work was supported by Canadian Institutes of Health Research Doctoral Award DFS- 157838 to M.R.W., National Institutes of Health R01 NS108484 and R01 EB011556 to A.K.D., R01 MH102595 and R01 MH117426 to M.R.D., and R37 NS056839 to T.A.J.. This work was performed with the support of the Mouse Genetic Engineering Facility (RRID:SCR 021927), a core facility within the Center for Biomedical Research Support at the University of Texas at Austin.

## Supplemental figures and tables

**Supplemental Figure 1.**
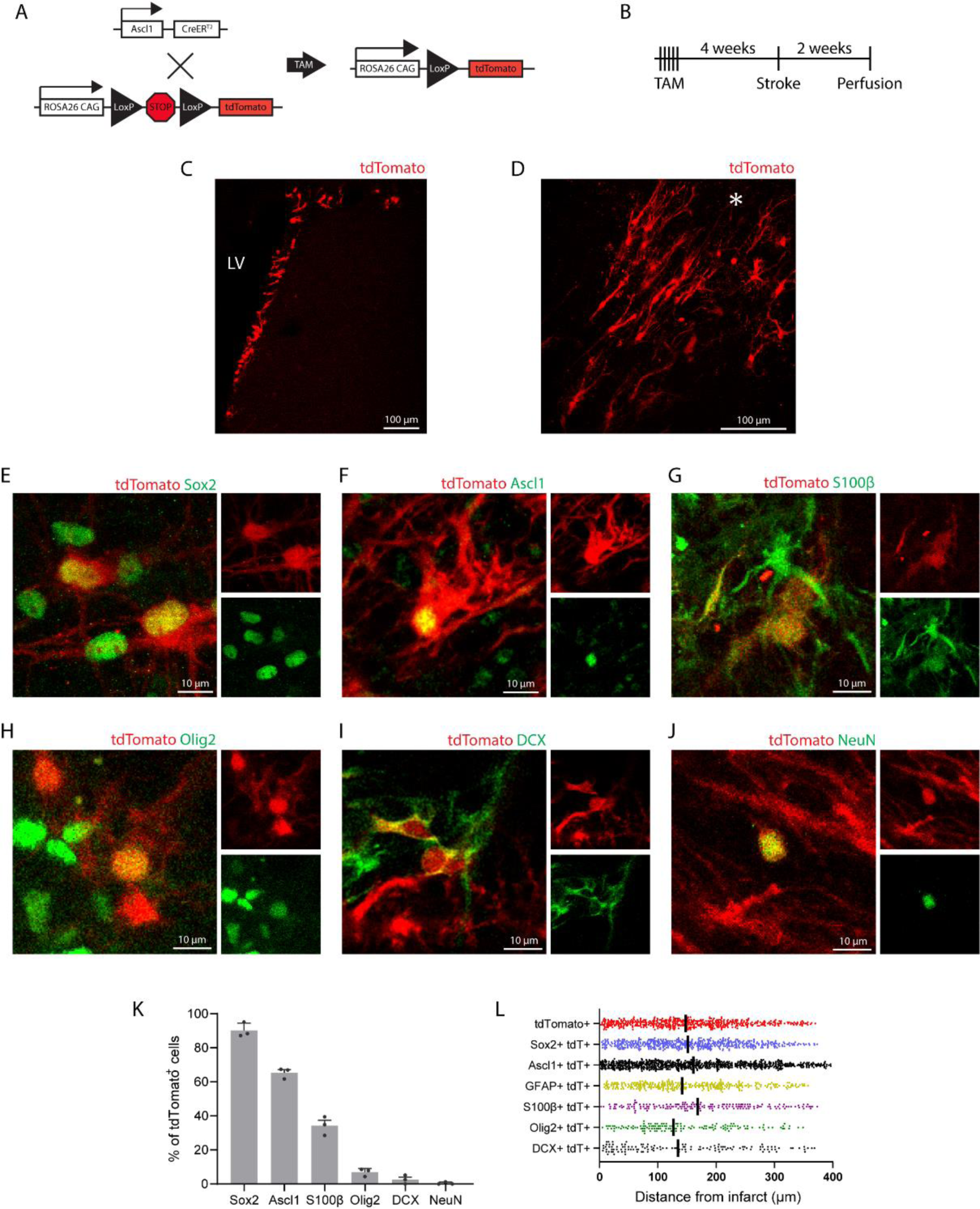
SVZ-derived cells are predominantly undifferentiated precursors and astrocytes. A) Schematic of genetic lineage tracing system where tamoxifen (TAM) induces indelible expression of tdTomato in Ascl1-expressing neural stem cells, intermediate progenitor cells, and their progeny. B) Experimental design. C) Image showing tdTomato expression in the SVZ. LV, lateral ventricle. D) Representative image of tdTomato^+^ cells localized to peri-infarct cortex 2 weeks after a cortical stroke (asterisk). E-J) Lineage tracing using Ascl1-CreER; Ai14 mice corroborates the major finding of Figure 1, that SVZ-derived cells in peri-infarct cortex are predominantly undifferentiated precursors and astrocytes. Representative images illustrating co-labeling of lineage traced tdTomato^+^ cells with differentiation stage-specific markers at two weeks post-stroke (E) Sox2, F) Ascl1, G) S100β, H) Olig2, I) DCX. J) NeuN). K) Quantification of marker expression by % of tdTomato^+^ cells. Data are presented as mean ± SEM. L) Spatial distribution of cells relative to the infarct border by marker expression at 2 weeks post-stroke in Nestin-CreER; Ai14 mice. Each point indicates a single cell. Vertical lines indicate medians. There was no clear pattern of spatial organization by cell type. Distance was measured as the distance to the nearest part of the infarct border within a given section for each cell. Only cells within 400 µm of the infarct border and within cortex were included in this analysis.

**Supplemental Figure 2.**
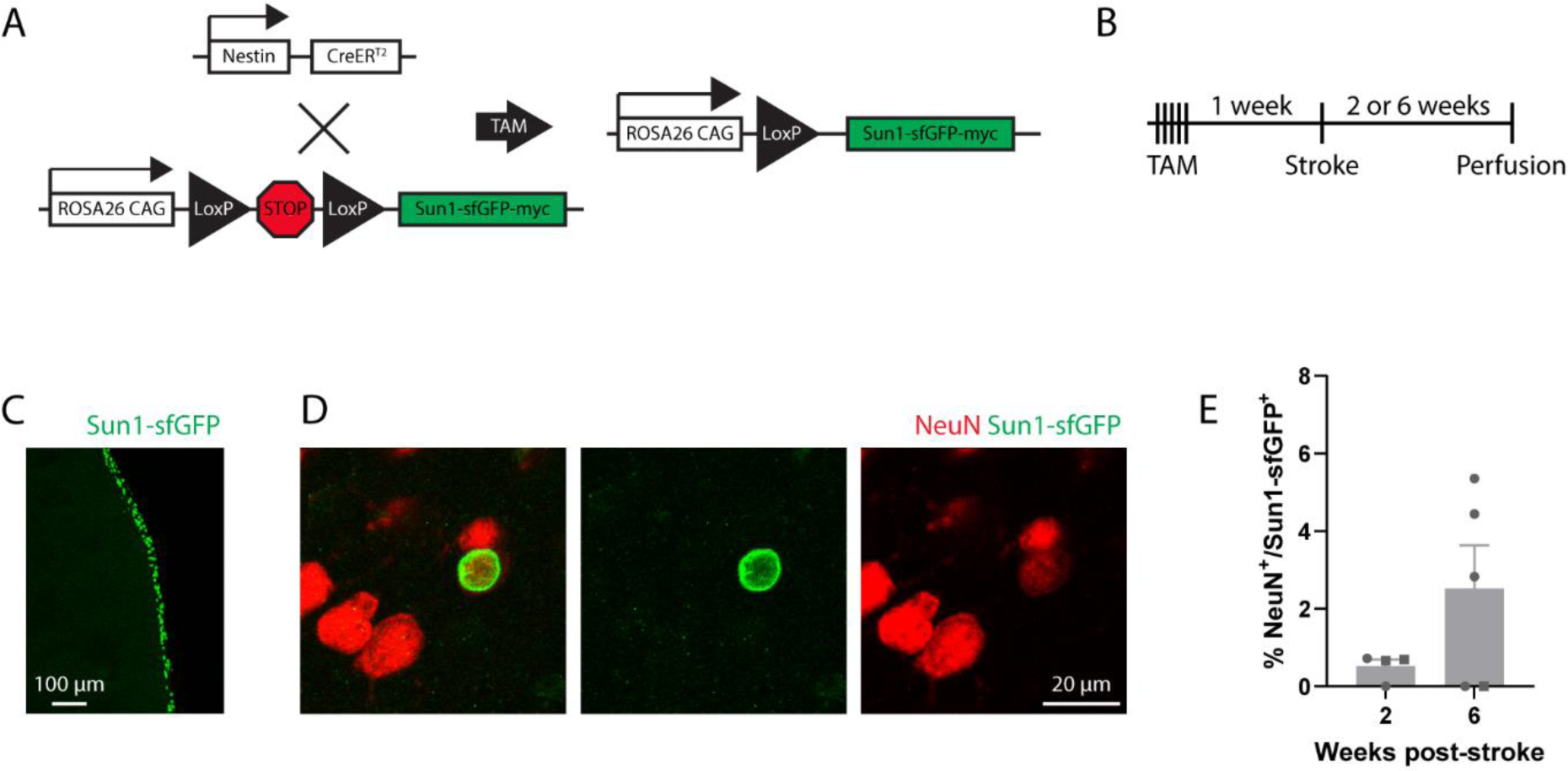
Few SVZ-derived cells become neurons. A) Schematic of genetic lineage tracing system where tamoxifen (TAM) induces indelible expression of a nuclear membrane-bound Sun1-sfGFP in neural stem cells and their progeny. B) Experimental design. C) Image showing Sun1-sfGFP expression in the SVZ. D) Representative image of a NeuN^+^Sun1-sfGFP^+^ nucleus in peri-infarct cortex. E) Quantification of SVZ-derived neurons. Data were derived from a combined 1014 nuclei counted across nine mice (n = 4 at two weeks, n = 5 at six weeks). There was not a significant difference between time points (t(4.2) = 1.79, p = 0.145, Welch’s corrected t test). Data are presented as mean ± SEM. Datapoints representing males are shown as circles; datapoints representing females are shown as squares.

**Supplemental Figure 3.**
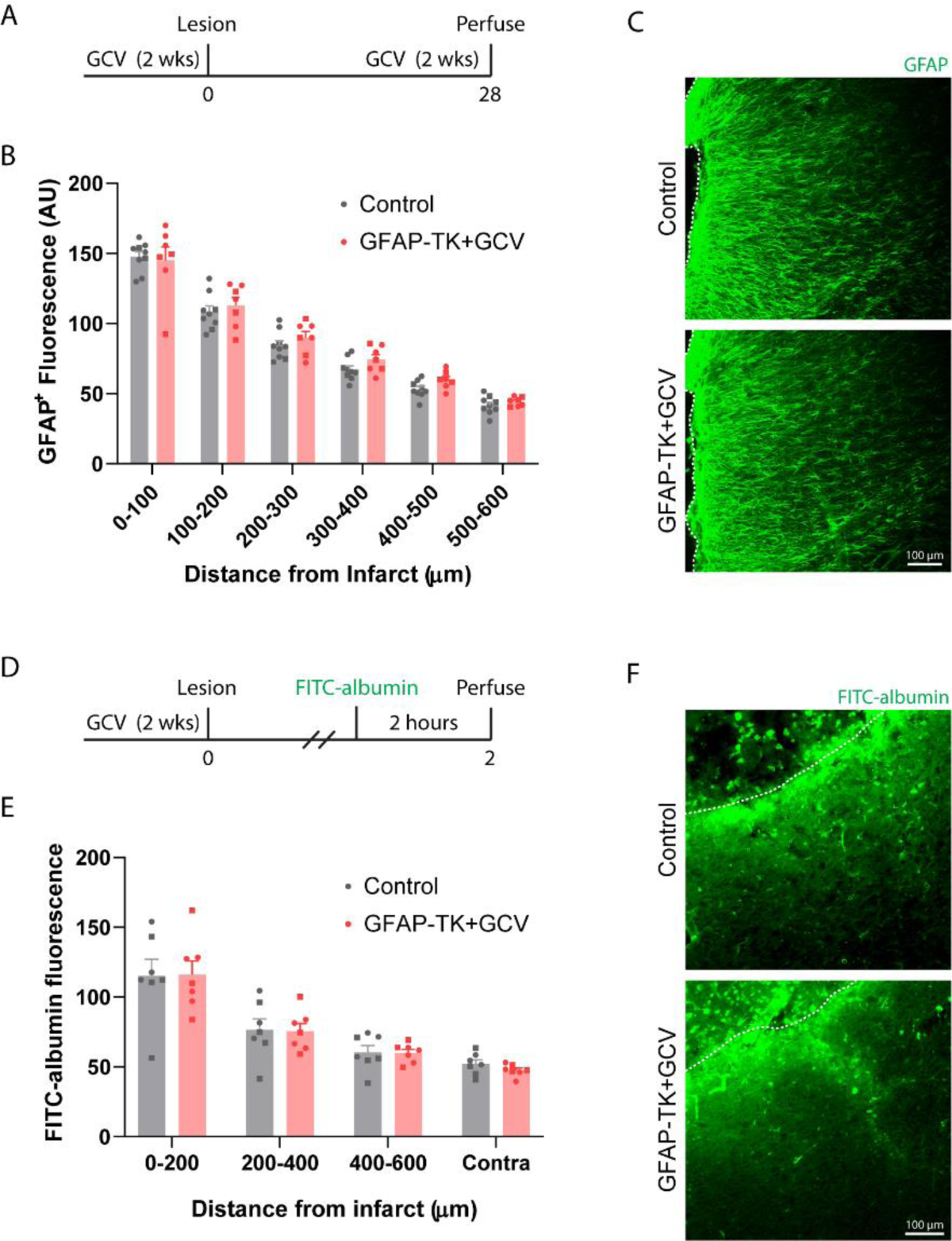
Neural stem cell ablation does not alter parenchymal astrocyte reactivity or vascular permeability after stroke. A) Experimental timeline for examining GFAP fluorescence. Tissue was obtained 28 days post-stroke (n = 9 controls (TK^-/-^), n = 7 GFAP-TK^+/-^). To assess parenchymal astrocyte reactivity, images were taken in superficial peri-infarct cortex where few SVZ-derived cells localize. B) GFAP^+^ fluorescence was increased near the lesion, but declined with distance away, characteristic of astrocyte reactivity (distance effect F(5, 84) = 172.9, p < 0.001). There was no significant group effect (F(1, 84) = 2.8, p = 0.100). C) Representative images of GFAP immunostaining in peri-infarct cortex. Dashed lines indicate the lesion border. D) Experimental design for examining vascular permeability. Two days post-stroke, animals were retro-orbitally injected with FITC-conjugated albumin 2 hours before perfusion (n = 7 controls (TK^-/-^), n = 7 GFAP-TK^+/-^). E) FITC-albumin fluorescence was greatest near the infarct border and declined with distance away, consistent with injury-induced vascular permeability (distance effect F(3, 48) = 36.9, p < 0.001). There was no significant effect of group (F(1, 48) = 0.07, p = 0.789). F) Representative images of FITC-albumin fluorescence in peri-infarct cortex. Dashed lines indicate the lesion border. Data are presented as mean ± SEM. Datapoints representing males are shown as circles; datapoints representing females are shown as squares.

**Supplemental Figure 4.**
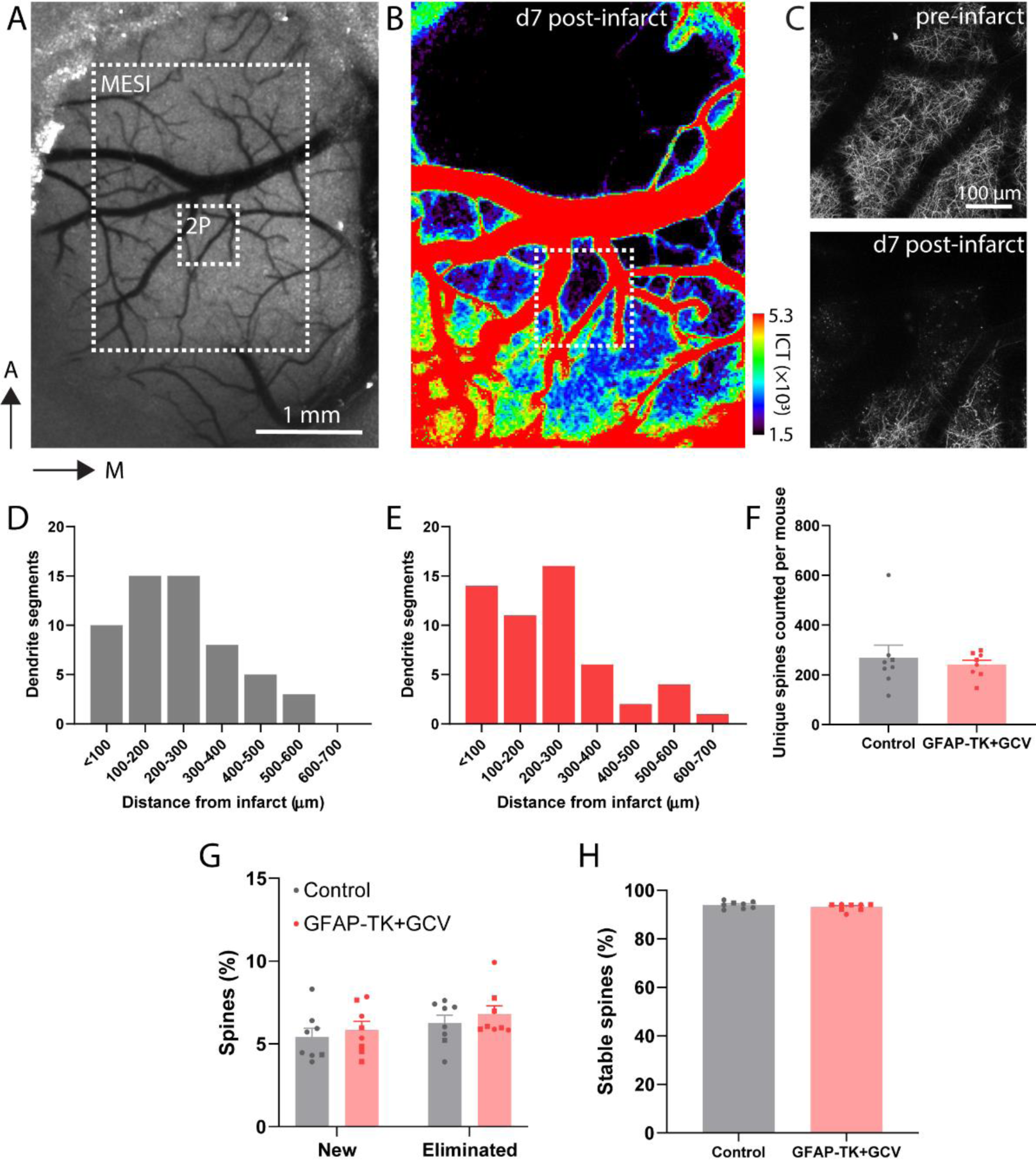
Additional data 2-photon imaging data. A) MESI accurately delineates the infarct border. Widefield laser speckle contrast image showing the cortical surface through a cranial window. B) Multi-exposure speckle imaging (MESI) image of blood flow corresponding to the region indicated in panel A. C) Two photon image of GFP-labeled apical dendrites (Thy1-GFP) corresponding to the region indicated in panels A and B. The infarct border revealed by MESI (black region in B) matches the region in which GFP fluorescence is absent. D-E) Distribution of the locations of analyzed dendrite segments relative to the infarct border for control (D) and GFAP-TK+GCV mice (E) (number of segments is summed across mice). F) Numbers of unique longitudinally tracked dendritic spines by group. Each point corresponds to an individual animal. Pre-stroke spine turnover (G) and stability (H) were not different between groups (t(14) ≤ 1.1, p ≥ 0.272). Data are presented as mean ± SEM. Datapoints representing males are shown as circles; datapoints representing females are shown as squares.

**Figure S5.**
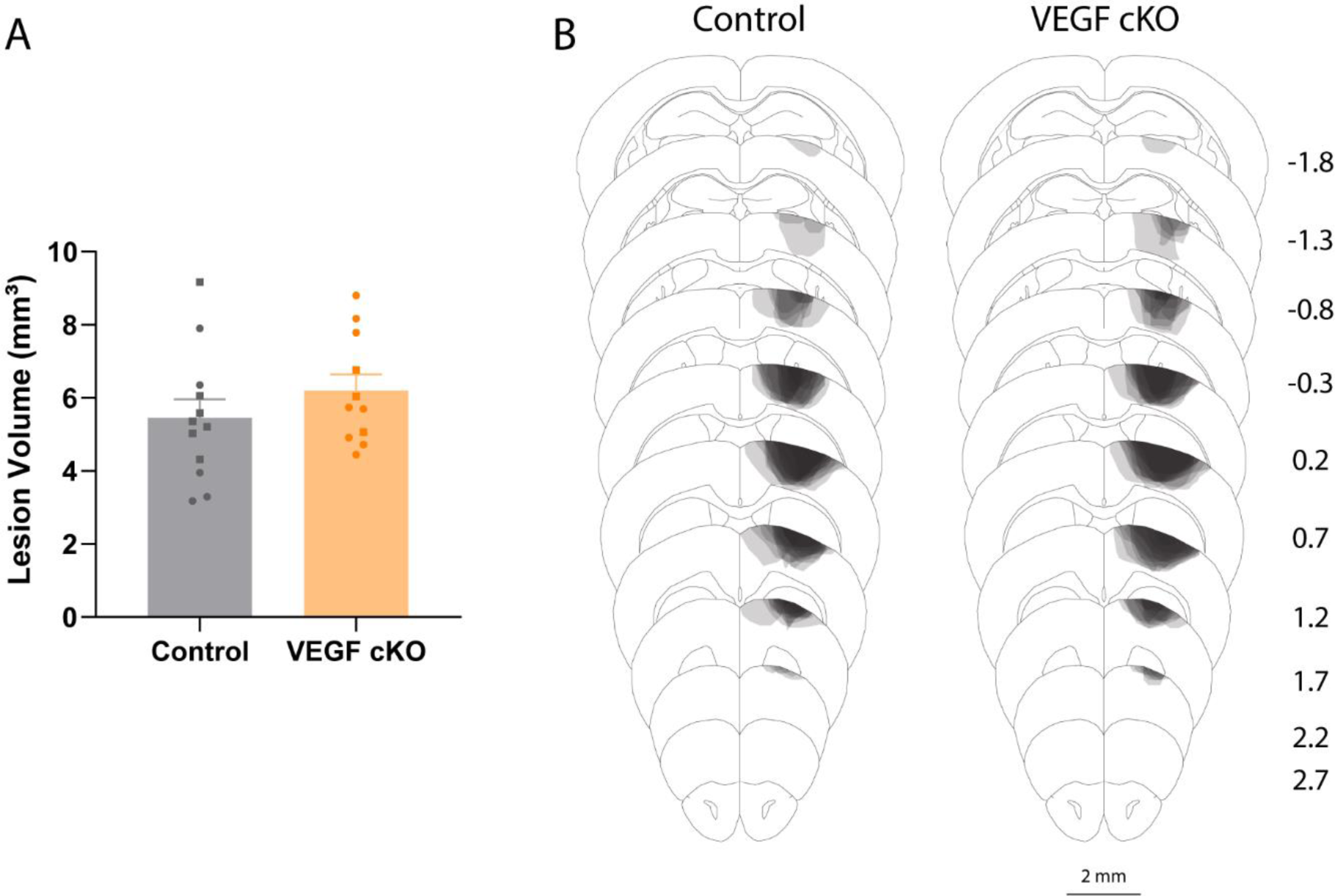
VEGF cKO did not affect lesion size or location. A) Lesion size was not different between groups (t(21) = 1.08, p = 0.292). B) Lesion reconstruction. Darker shades indicate greater overlap between animals.

**Figure S6.**
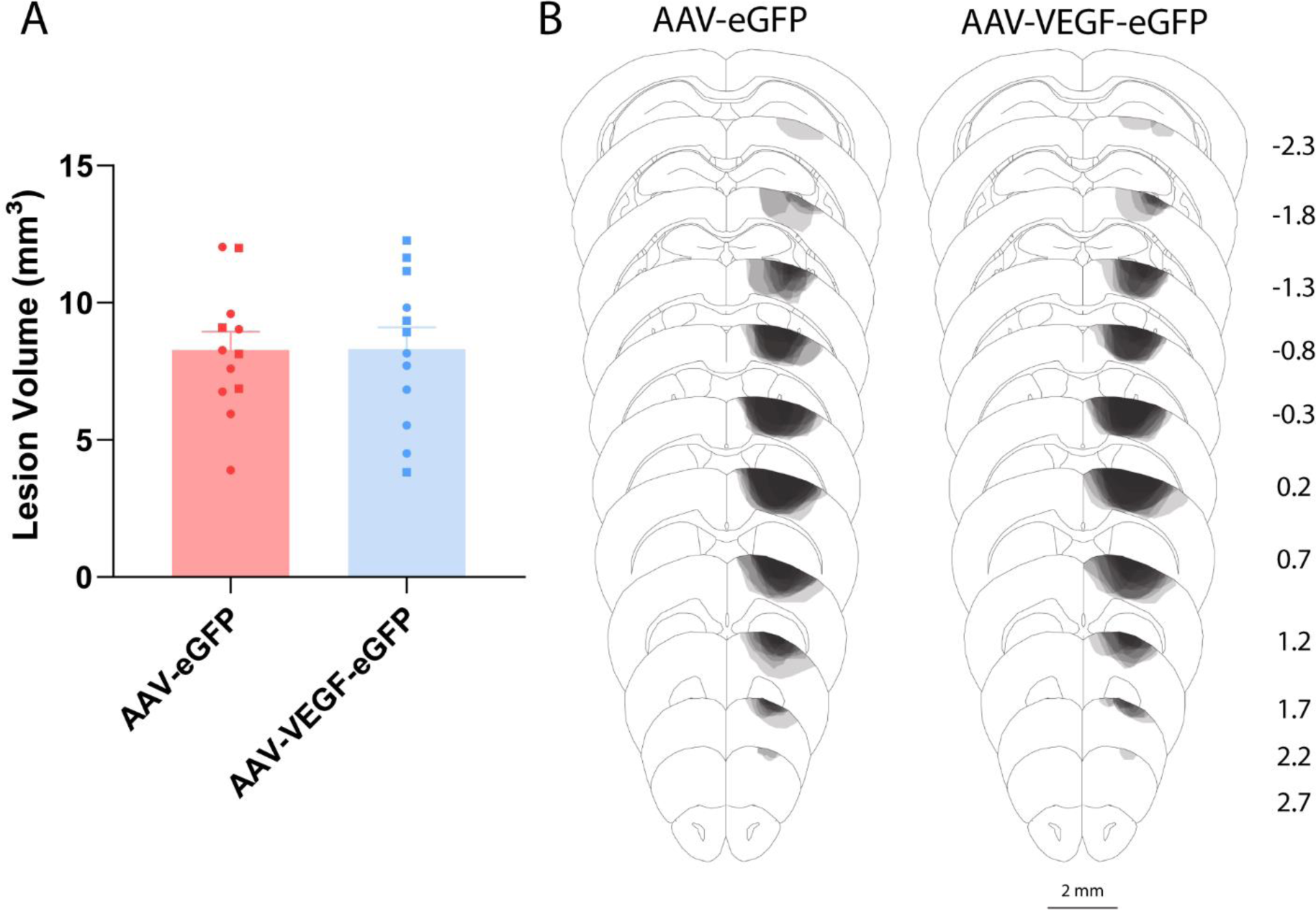
AAV-VEGF-eGFP did not affect lesion size or location. A) Lesion size was not different between groups (t(22) = 0.04, p = 0.967). B) Lesion reconstruction. Darker shades indicate greater overlap between animals.

**Figure S7.**
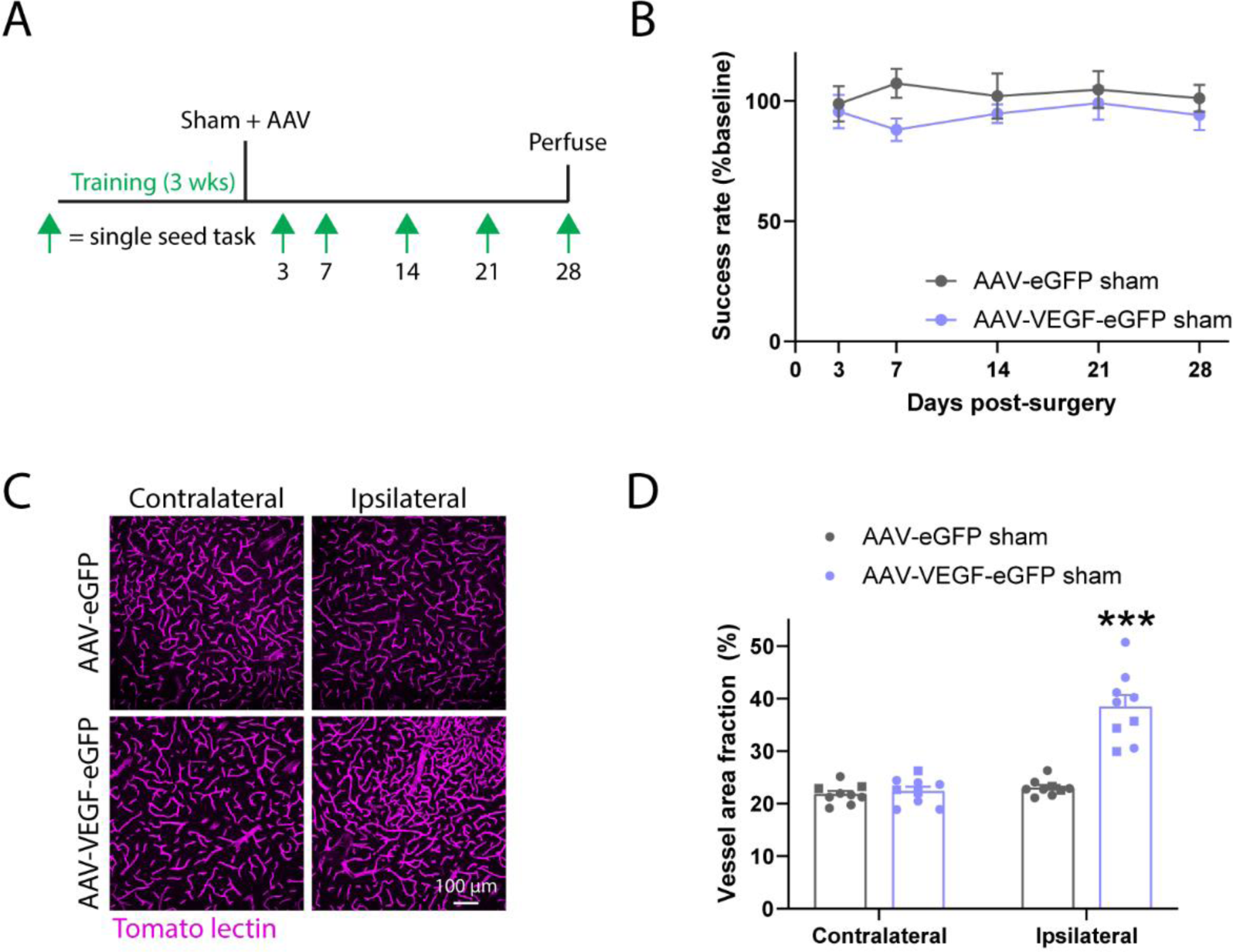
AAV-VEGF-eGFP does not affect motor function in sham-operated animals. A) Experimental timeline. Wildtype mice were trained on the single seed reaching task, subjected to a sham stroke procedure, and injected with either AAV-eGFP or AAV-VEGF-eGFP (n = 8/group). Performance on the single seed task was periodically tested up to 28 days post-surgery. B) Performance on the single seed task was not different between groups (F(1, 14) = 3.8, p = 0.071). C, D) Representative images (C) and quantification (D) of vasculature in contralateral and ipsilateral cortex (relative to AAV injection site). AAV-VEGF-eGFP increased vessel density in ipsilateral cortex (t(16) = 6.8, ***p < 0.001) (n = 9/group). Data are presented as mean ± SEM. Where individual datapoints are shown, datapoints representing males are shown as circles; datapoints representing females are shown as squares.

**Supplemental Table 1.**
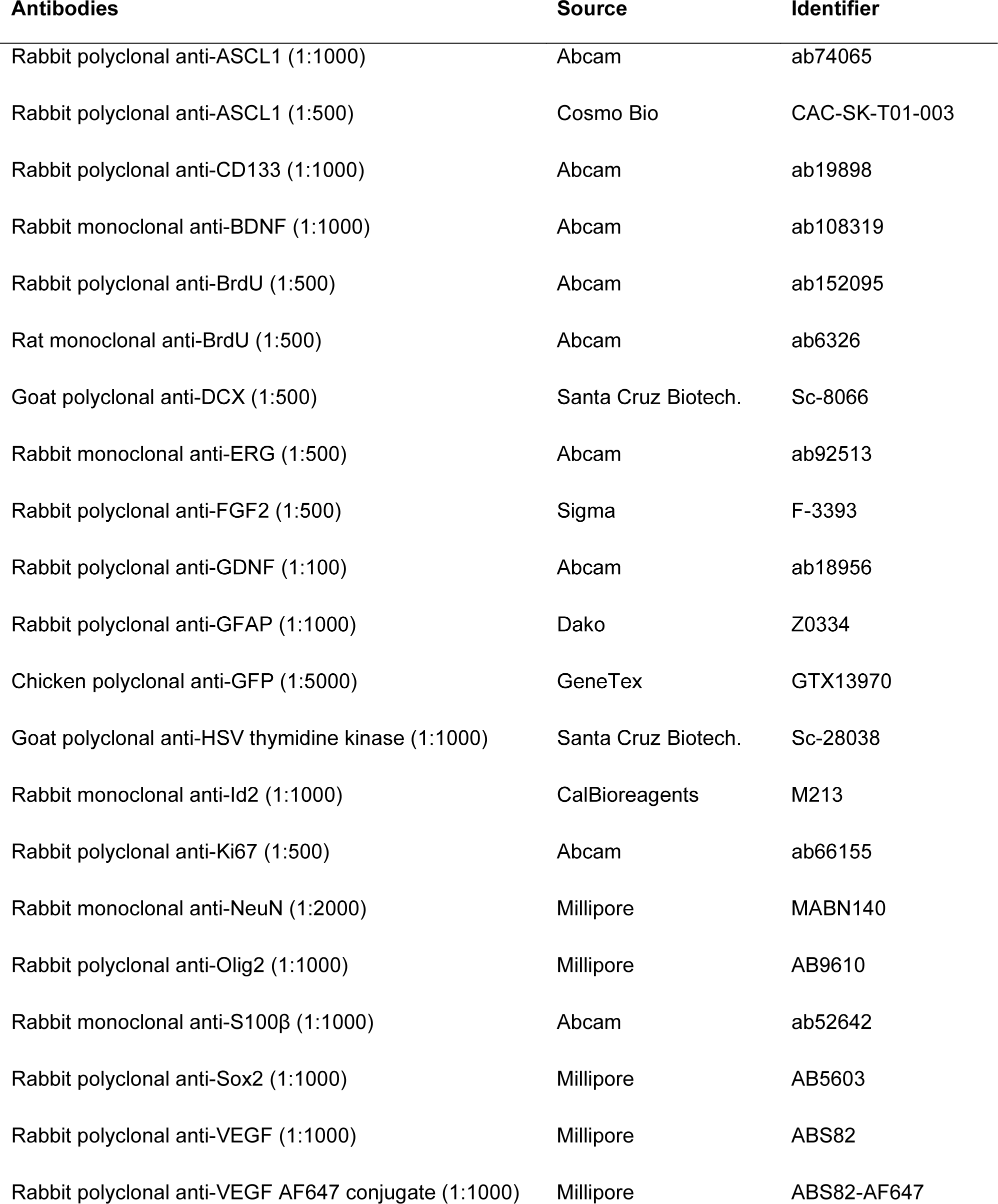
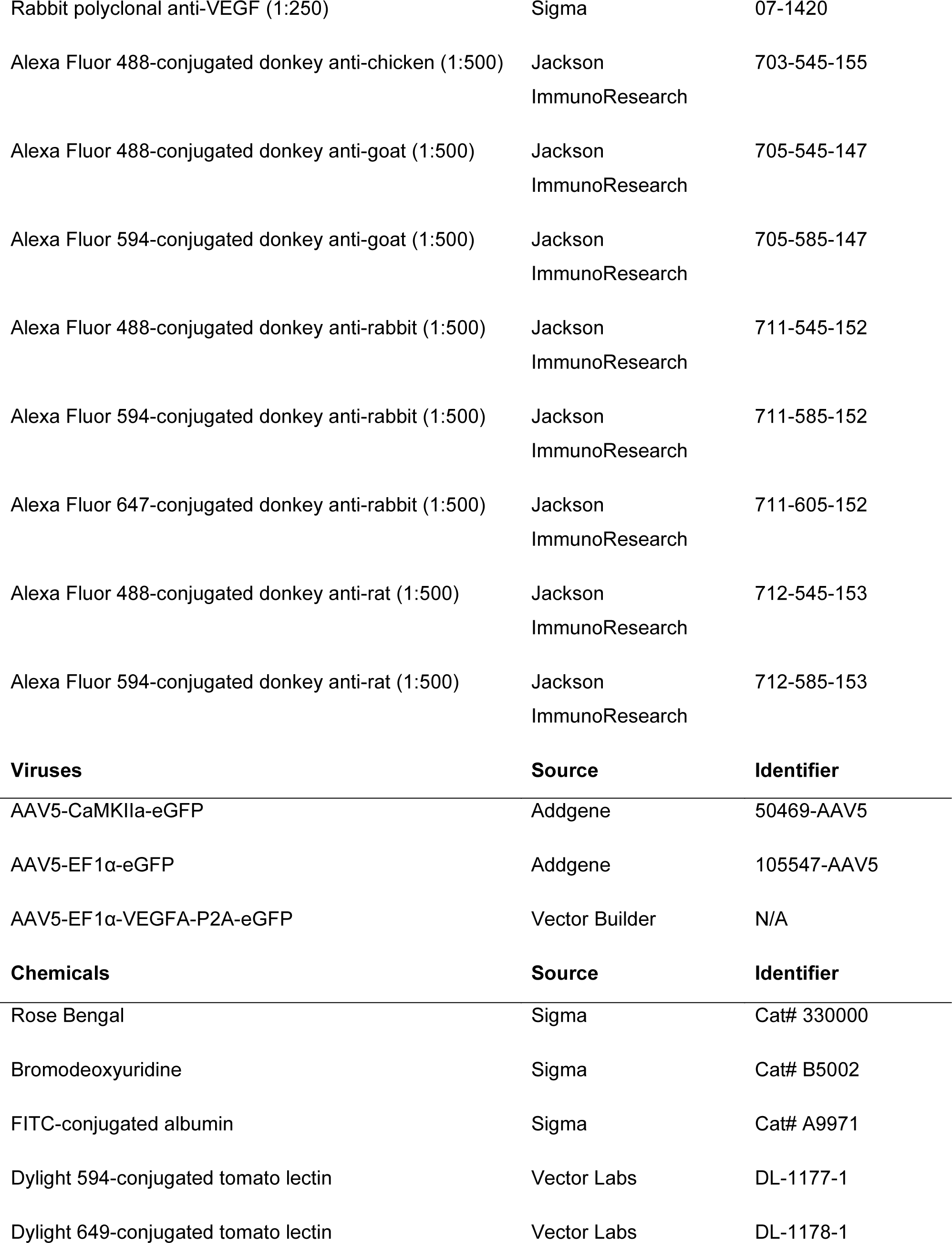
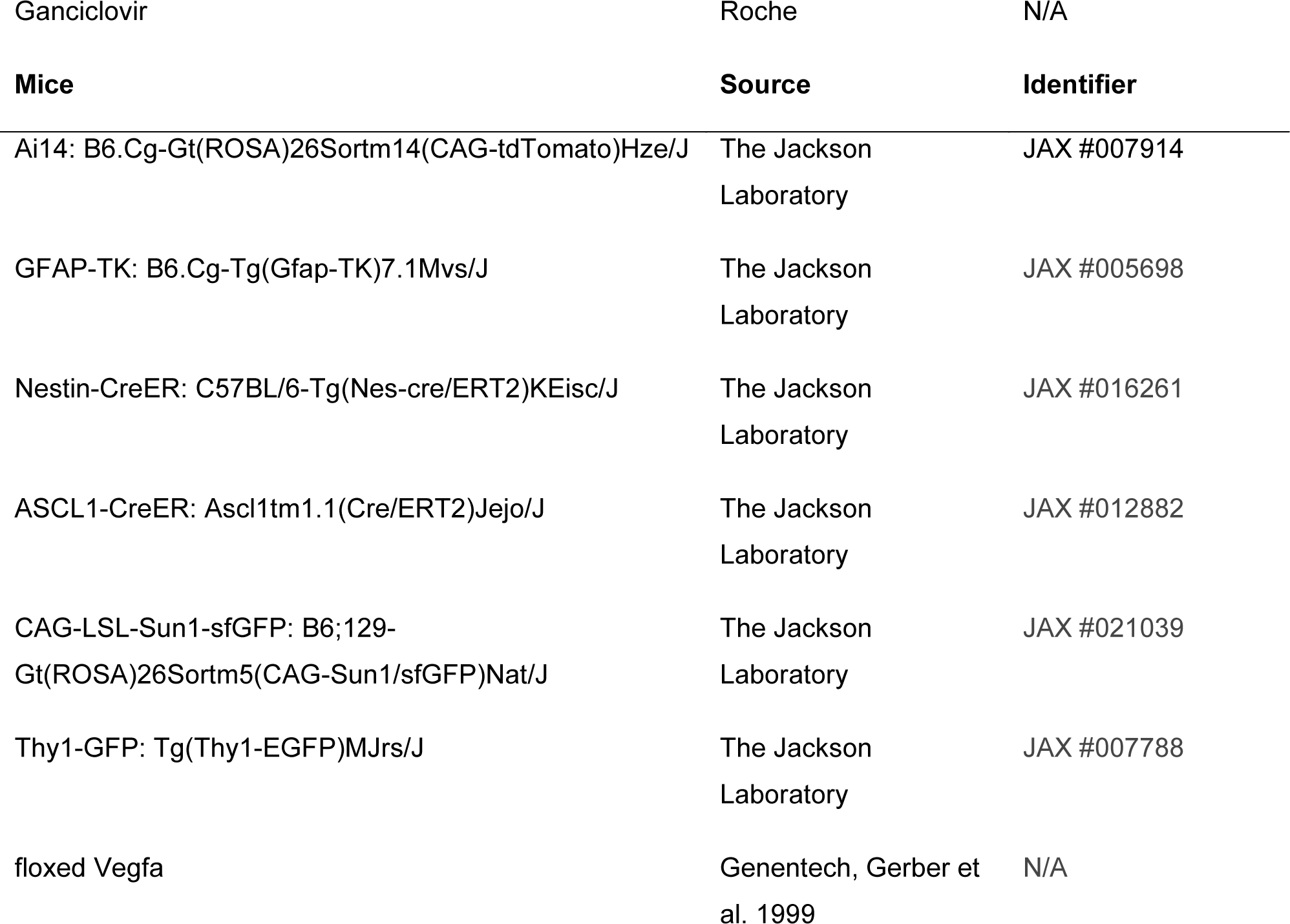
List of reagents.

